# Automated Feature Extraction from Population Wearable Device Data Identified Novel Loci Associated with Sleep and Circadian Rhythms

**DOI:** 10.1101/2020.03.31.017608

**Authors:** Xinyue Li, Hongyu Zhao

## Abstract

Wearable devices have been increasingly used in research to provide continuous physical activity monitoring, but how to effectively extract features remains challenging for researchers. To analyze the generated actigraphy data in large-scale population studies, we developed computationally efficient methods to derive sleep and activity features through a Hidden Markov Model-based sleep/wake identification algorithm, and circadian rhythm features through a Penalized Multi-band Learning approach adapted from machine learning. Unsupervised feature extraction is useful when labeled data are unavailable, especially in large-scale population studies. We applied these two methods to the UK Biobank wearable device data and used the derived sleep and circadian features as phenotypes in genome-wide association studies. We identified 53 genetic loci with p<5×10^-8^ including genes known to be associated with sleep disorders and circadian rhythms as well as novel loci associated with Body Mass Index, mental diseases and neurological disorders, which suggest shared genetic factors of sleep and circadian rhythms with physical and mental health. Further cross-tissue enrichment analysis highlights the important role of the central nervous system and the shared genetic architecture with metabolism-related traits and the metabolic system. Our study demonstrates the effectiveness of our unsupervised methods for wearable device data when additional training data cannot be easily acquired, and our study further expands the application of wearable devices in population studies and genetic studies to provide novel biological insights.

## Introduction

Sleep is essential for human health and well-being, and changes in sleeping patterns or habits can negatively affect health, leading to physical and mental disorders^1–4^. The corresponding sleepwake circadian rhythm is also essential for human health. Circadian rhythms are endogenous biological processes that follow a period of approximately 24 hours and are entrained by environmental stimuli such as the light/dark cycle to adjust the 24-hour cycle^5^. The circadian system is important for sleep regulation, and dysregulated sleep-wake circadian rhythms can cause diseases including sleep disorders, metabolic syndrome, and psychiatric and neurodegenerative diseases^6–8^. While it is vital to obtain a thorough understanding of the important roles of sleep and circadian rhythms in human health, they still remain poorly understood.

Actigraphy has been increasingly used in sleep and circadian studies, as it can provide continuous and objective activity monitoring and is low-cost and easy to wear. Actigraphy can address some of the limitations with traditional sleep diaries, including subjectivity, bias, difficulty in completion by young children or patients, and extra manual work for caregivers. While polysomnography (PSG) as the “gold standard” in sleep studies does not have the same issues as sleep logs, it is limited by high costs, in-lab setting, intrusive measures, and difficulty in long-time monitoring. The continuous and objective measures provided by actigraphy can provide reliable information on sleep^9^, and it is especially useful for studying long-duration circadian rhythms. In cases of large-scale epidemiological studies, the availability of actigraphy data provides excellent opportunities for studying population-level and individual-level sleep characteristics and circadian rhythm patterns. However, the analysis of actigraphy data remains a major obstacle for researchers.

One major challenge in the application of actigraphy is to extract sleep and circadian rhythm features without additional information, such as sleep diaries and/or PSG validation that are often unavailable or labor-intensive to collect. To extract sleep features such as sleep start, sleep end, and sleep duration, it is typical to either obtain the gold-standard PSG records or obtain sleep dairies on go-to-sleep time and wake-up time to build sleep identification algorithms^10–13^. To ease researcher’s efforts in collecting additional data and still accurately infer sleep features, there is a need to develop necessary methodology to infer sleep parameters based on actigraphy in the absence of sleep records. The method will be useful in circadian studies where PSG for long-time monitoring cannot be acquired and continuous activity logs requires much manual work. It is also particularly useful in large-scale epidemiological studies where collecting PSG for all participants is unrealistic and recording sleep logs is labor-intensive.

In this paper, we analyzed data from the UK Biobank study, where accelerometer data from over 100,000 participants and genetic data from near 500,000 participants are available. We applied novel data processing methods to the accelerometer data from 90,515 participants after quality control procedures to extract sleep and circadian rhythm features. We then conducted genome-wide association studies (GWAS) and identified 19 and 34 genetic loci associated with sleep traits and circadian rhythm traits at p<5×10^-8^ respectively, of which 5 and 13 loci reached the significance level p<5×10^-9^. Further tissue enrichment analysis highlights the important roles of the central nervous system and the metabolic system, thereby providing new insights into the molecular regulation and genetic basis of sleep and circadian rhythms.

## Results

### Loci Associated with Sleep-Activity Traits and Circadian Traits

The Manhattan plots of GWAS results for HMM inferred sleep and activity traits are shown in Supplementary Figure S1. The heritability estimates from LD score regression^14^ for mean activity levels during sleep and during wake are ~5% and ~7%, respectively. For mean activity levels during sleep, four independent regions on chromosomes 2, 5, 6, and 14 contained significant SNPs with p-value < 5×10^-8^, two of which had p-values < 5×10^-9^. The strongest association signal was on chromosome 2 at SNPs within gene MEIS1, known for association with Restless Leg Syndrome and Insomnia (Table 1).^15–18^ The other three genetic loci were rs188904275 in JAKMIP2 (p-value = 3.7×10^-8^), rs184670665 near IMPG1 (p-value = 2.4×10^-10^), and rs73586669 near OR4E1 (p-value = 2.4×10^-8^), with JAKMIP2 previously found to be associated with Body Mass Index (BMI) measurements and pulmonary diseases^19^. These novel loci were not previously associated with activity levels during sleep. SNP rs7087063 in gene CELF2 on chromosome 10 was found to be associated with the HMM estimated activity variability during wake (p-value = 3.0×10^-8^), and we note that CELF2 was previously found to be associated with Alzheimer’s disease^20^ (Table 1). No association was detected for HMM estimated activity variability during sleep or mean activity levels during wake. The QQ-plots checking for population stratification indicate that the population structure was properly controlled for, with small values of the inflation factor λ under 1.04 (Supplementary Figure S2).

**Table 1.**
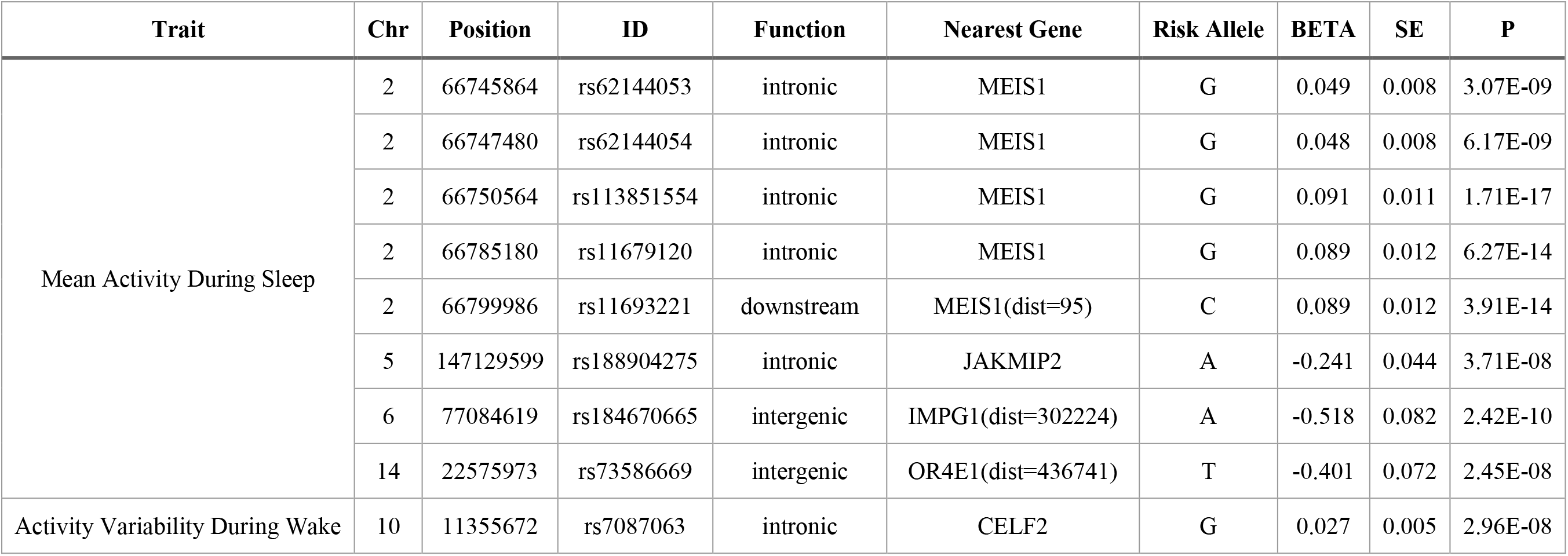
The SNPs identified in genome-wide association studies at the significance level of 5×10^-8^ that are associated with sleep and activity traits inferred from accelerometer-measured physical activity in 90,515 UK Biobank participants.

**Table 2.** The SNPs identified in genome-wide association studies at the significance level of 5×10^8^ that are associated with sleep duration traits inferred from accelerometer-measured physical activity in 90,515 UK Biobank participants.

| Trait | Chr | Position | ID | Function | Nearest Gene | Risk Allele | BETA | SE | P |
| --- | --- | --- | --- | --- | --- | --- | --- | --- | --- |
| Sleep Duration < 5h | 7 | 139916399 | rs573901234 | intergenic | JHDM1D-AS1(dist=36959) | T | 3.294 | 0.210 | 1.44E-08 |
| Sleep Duration > 10h | 1 | 32558622 | rs74460673 | intronic | TMEM39B | A | 3.822 | 0.242 | 2.96E-08 |
|  | 2 | 171183777 | rs573982927 | intronic | MYO3B | T | 3.460 | 0.227 | 4.45E-08 |
|  | 4 | 139129025 | rs182651559 | intronic | SLC7A11 | T | 4.989 | 0.280 | 9.72E-09 |
|  | 8 | 131387296 | rs571444813 | intronic | ASAP1 | T | 5.568 | 0.314 | 4.62E-08 |
|  | 22 | 24204438 | rs138381486 | intronic | SLC2A11 | T | 1.797 | 0.107 | 4.54E-08 |

The Manhattan plots of GWAS results for sleep duration, sleep start and sleep end traits are shown in Supplementary Figure S1. The heritability estimates from LD score regression^14^ for sleep start and sleep end are ~8% and ~7%, respectively. SNP rs573901234 near gene JHDM1D-AS1 on chromosome 7 was associated with short sleep duration < 5 hours (p-value = 1.4×10^-8^). Five SNPs were associated with long sleep duration > 10 hours, among which rs74460673 at gene TMEM39B was previously found to be associated with BMI^21, 22^ and rs573982927 at MYO3B was associated with obesity traits^23, 24^. Twenty seven SNPs were found to be associated with sleep start, including three SNPs at or near gene MEIS1 related to Restless Leg Syndrome and insomnia^15–18^ and nineteen SNPs at gene BTBD9 also related to Restless Leg Syndrome^25^ (Supplementary Table S1). Sleep start is also associated with one SNP at gene CYP7B1 related to BMI, two SNPs near gene HSD17B12 related to BMI^21, 26, 27^, two SNPs near MIR129-2 also related to BMI^21, 22^ and Alzheimer’s Disease^28^, and two SNPs near LOC101928944 related to schizophrenia^29, 30^. As for sleep end, association signals were found for five SNPs in the intergenic regions near LINC02260, which is known to be related to red blood cell measures such as cell counts and hemoglobin content^31^, and blood cell information is also known to be associated with sleep deprivation and sleep disorders^32–34^. Sleep end is also associated with one SNP at gene NTNG1 related to Restless Leg Syndrome^25^ and BMI^21, 35^, one SNP near LINC00963, and one SNP near GLRX3.

From the penalized multi-band learning approach, the most dominant periodicities are: 1-day, 1/2-day, and 1/3-day. The Manhattan plots for the GWAS results are shown in Supplementary Figure S1. The heritability estimate from LD score regression^14^ for the circadian feature 1-day periodicity is 9%. For the strength of 1-day periodicity, five circadian SNPs were identified: rs189005747 at gene XKR4 (p-value=9.9×10^-9^), rs534035399 at LINC01508 (p-value=1.8×10^-9^), rs144874087 near LINC01935 (p-value=1.9×10^-8^), rs181820530 near LINC01935 (p-value=4.5×10^-8^), and rs554696049 at LINC01501 (p-value=4.8×10^-8^), in which XKR4 was previously found to be associated with thyroid stimulating hormone^36–38^ and coronary artery disease, and LINC01508 and LINC01501 are RNA genes (shown in Supplementary Table S2).

1/2-day periodicity measures the strength of day-night rhythmicity^39^ and the strongest association signals (p-value < 5×10^-9^) were detected in the intergenic regions near UBE2F-SCLY and FBXO15 and in the intronic regions at FYB1 and CFAP44, in which intergenic regions near FBXO15 were previously found to be associated with insomnia^40^ and FYB1 was associated with depression^41, 42^. There were also association signals for SNP clusters in the intergenic regions near gene TNR as well as for SNPs at RASEF, TMEM132D, ERCC2, MPPED1 and near BRINP3, LINC01287, and MLYCD (p-value < 5×10^-8^; Supplementary Table S2).

1/3-day periodicity measures the 1/3-day rhythmicity that not only involves activities during the day but also captures activities during sleep^39^. The strongest signals (p-value < 5×10^-9^) were identified at SNP clusters in the intergenic regions near BRINP3, URB2, GRIA1 and LOC400682 and in the intronic regions at MGAT5, C3orf20, and LINC01861, where BRINP3 was associated with BMI measurement^21, 22, 27^, depression^43^, and rheumatoid arthritis^44^, and GRIA1 was associated with schizophrenia^29, 45, 46^ and circadian rhythm^47^. A cluster of five SNPs at CDH6 reached significance level at 5×10^-8^, and CDH6 was associated with resting heart rates^48^. Details on other novel SNP associations are listed in Supplementary Table S2. QQ-plots show that the estimated values of the inflation factor λ are under 1.07 and that the population structure was properly controlled for (Supplementary Figure S2).

With respect to the quality control procedures for all GWAS, the inflation factor λ was under 1.12 for all traits considered, suggesting appropriate control of population structure in the analysis (Supplementary Figure S2). LD Score intercepts estimated using LD score regression^14^ were in the range of 0.99-1.02, indicating that the genomic inflation was due to polygenic architectures rather than uncorrected population structure. Further, we estimated partitioned heritability^49^ using ten broad tissue categories and no tissue was significantly enriched. For all sleep time-related traits, the adrenal/pancreatic tissues were relatively more enriched than the other tissues (Supplementary Figure S3). For activity levels during sleep, the cardiovascular tissues were relatively more enriched compared to the other tissues (Supplementary Figure S3).

### Genetic Correlation of Sleep and Daily Rhythm with Other Traits

To further investigate the relationships of sleep and circadian traits with other complex traits, we estimated genetic correlation with a number of traits, including screen exposure, sleep, mental health, BMI and diet, alcohol consumption, shift-work, and diseases such as respiratory diseases and anemia (Supplementary Table S3) and we set the statistical significance threshold at 7.9 × 10^-5^ through Bonferroni correction (12 sleep and circadian traits × 53 traits). For sedentary and screen exposure traits, we observed significant negative genetic correlation of time spent watching TV or using computer with the strength of circadian rhythmicity (correlation=-0.300, p=2.1 × 10^-12^ and correlation=-0.294, *p* =5.7 × 10^-10^, respectively), and we also observed significant negative correlation of time spent using computer with sleep duration, sleep start (go to bed early), and activity levels during the wake status(correlation=-0.238, *p* =2.7 × 10^-7^; correlation=-0.353, *p* =3.5 × 10^-5^ and correlation=-0.200, *p* =1.8 × 10^-22^ respectively). There was positive genetic correlation between the time spent watching TV and activity levels during sleep (correlation=0.203, *p* =7.3 × 10^-5^). For BMI and diet related traits, we observed significant negative genetic correlation between BMI and weight with the strength of circadian rhythm (1-day periodicity), activity levels when awake, and sleep duration (all negative correlations with p-values < 7.9 × 10^-5^). We did not observe significant genetic correlation with mental health traits, alcohol consumption, shift-work, and respiratory diseases either. While the genetic correlation with anemia was not significant, there was moderate positive genetic correlation between activity levels during sleep and iron deficiency anemia (correlation=0.404, *p* =6.8 × 10^-3^), and iron deficiency is known to be associated with poor sleep quality^68^ and restless legs syndrome^50, 51^.

We also observed significant positive genetic correlation between accelerometer-derived and self-reported sleep duration, between accelerometer-derived sleep start/end and self-reported chronotypes, and between accelerometer-derived sleep end and self-reported hypersomnia (all p-values<7.9× 10^-5^). These results suggest agreement between accelerometer-derived measures and self-reported measures for sleep time and are also consistent with previous studies^52^. We did not observe significant genetic correlation of accelerometer-derived measures with sleep disorders, possibly because the ambiguity in the definition of sleep disorders. We did not observe significant genetic correlation between accelerometer-derived activity level measures and self-reported physical activity measures.

For the significant trait-pair of BMI and 1-day periodicity denoting the strength of circadian rhythm, we further conducted two-sample MR analysis using GWAS summary statistics from the GIANT study^27^. The estimated causal effects across different MR estimation methods are negative for both directions but were not statistically significant.

### Cross-Tissue Transcriptome-Wide Association Analysis

We applied UTMOST^53^ to perform tissue enrichment analysis in 44 tissue types and identified single-tissue and cross-tissue gene-trait associations. For single-tissue tests, 124 gene-trait pairs were identified (Supplementary Table S4). There were 43 unique genes in total and 16 gene-trait pairs were identified in more than one tissue type (p-values < 3.3 × 10^-6^ after Bonferroni correction for 15,000 genes^53^). Among them, the GLTP-sleep duration pair was identified in 38 tissue types, which is a novel association not reported in previous studies, and GLTP is related to Glycolipid Transfer Protein for protein binding and lipid binding^54^. L3MBTL2, Lethal (3) malignant brain tumor-like protein 2 related to protein binding and DNA regulation^55^, was associated with more than one trait, including activity levels during sleep and wake as well as the circadian rhythm, and the three L3MBTL2-trait pairs were all significant in the subcutaneous adipose tissues. Most genes that appeared in more than one gene-trait pair have functions in protein binding, including CEP70, GIPC2, TRAF3, ABCD2, LIMS1, SEPN1. BRSK1 and LRFN4 are related to neurotransmitter activities^55^. TREH is related to digestion and galactose metabolism^55^, and the TREH-sleep start pairs are significant in mucosa esophagus, stomach, and transverse colon, all of which belong to the digestive system. Among all 44 tissue types, subcutaneous adipose, brain anterior cingulate cortex, and skeletal muscles are most enriched with seven genetrait pairs (Supplementary Figure S4). Tissue enrichment analysis through UTMOST identified novel genes associated with sleep and circadian traits and highlighted the relevance of tissues in the central nervous system and the metabolic system in sleep and circadian regulation.

Joint tests for gene-trait associations across tissues identified 34 gene-trait pairs with p-values < 3.3 × 10^-6^ after Bonferroni correction for 15,000 genes^53^ and 20 gene-trait pairs with p-values < 3.3× 10^-7^ if Bonferroni correction further adjusts for 10 traits tested (Supplementary Table S5). Three genes, CA7, DYNC1LI1, and ELMOD2, were associated with more than one trait. CA7 was associated with activity levels during sleep and activity levels during wake, and in previous studies its expression in brain was associated with neurological disorders^56^. DYNC1LI1 was associated with sleep duration and daily rhythmicity and its functions are RNA binding^57^ and protein binding^58^. ELMOD2 was associated with activity variability during wake, sleep duration, sleep end and daily rhythmicity and its function may be related to respiratory diseases^59^. Among genetrait pairs, GRIA1 was also identified in GWAS analysis and was associated with circadian rhythms and sleep traits in previous studies^40, 47^, and its functions involve amyloid-beta binding in hippocampal neurons and ionotropic neurotransmitter receptor activities^60^. Gene Ontology (GO) enrichment analysis using DAVID^61^ did not identify significant enrichment among different GO terms.

## Discussion

Using information related to sleep and activity inferred from the UK Biobank accelerometer data, we identified five genetic loci associated with HMM derived sleep related traits. The association of activities during sleep with restless leg syndrome, obesity, and pulmonary diseases may be attributed to the fact that individuals with these traits may be less likely to sleep well at night and are prone to being restless. Sleep start and end are also strongly associated with Restless Leg Syndrome, as people with this disorder may have difficulty falling asleep or staying asleep, and because symptoms worsen at night only with a short period in early morning that is symptom-free, people with Restless Leg Syndrome tend to have late sleep onset^62^. Sleep duration, sleep start and sleep end are all associated with BMI, consistent with the established association between sleep and obesity. As previous studies^63^ discussed how decreased sleep duration can elevate obesity risks and how sleep disorders can increase risks for chronic health conditions, our study also provides evidence of genetic correlations among sleep, metabolism and obesity. For sleep end, it is interesting to identify SNPs near LINC02260, which was previously found to be associated with red blood cell related measures^31^. It is known from previous studies that sleep deprivation and sleep disorders can lead to changes in metabolism with altered red blood cell measurements such as increased red blood cell counts^32–34^. Thus, our study also confirmed the important link between sleep and blood cell metabolism, and the complex relationship remains to be studied.

For circadian and daily rhythm analysis, we were able to identify 13 loci associated with periodic traits. The SNPs located at FBXO15 and GRIA1 were associated with circadian rhythms and sleep traits in previous studies^40, 47^, and the associations of SNPs in XKR4 and CDH6 with thyroid stimulating hormones and resting heart rates can be explained by the circadian oscillation natures of hormones and heart rates.^64, 65^ These association results suggest that the Penalized Multiband Learning approach^39^ in extracting clinically meaningful circadian features from objective physical activity measures. Furthermore, we identified novel associations for SNPs at or near FYB1, GRIA1, CDH6 and BRINP3, which are associated with BMI and mental problems, suggesting shared genetics between sleep-wake circadian rhythms and multiple traits.

Our study found that sleep and physical activity are closely related to mental and neurological problems, as the identified SNPs/loci from our GWAS results were also associated with traits including depression, schizophrenia and Alzheimer’s disease. Our cross-tissue transcriptome-wide association analysis also implied the important role of the central nervous system in physical activity, sleep, and circadian rhythm. Our results are consistent with previous UK Biobank studies^66, 67^ and confirmed the shared genetic architectures of sleep and physical activity with mental health and neurological disorders.

Our study shows strong evidence for shared genetics of activity and circadian rhythm with metabolism-related traits and the metabolic system. In addition to GWAS results that suggest shared genetic architecture with BMI, genome-wide genetic correlation estimates also provide strong evidence that a higher BMI is associated with lower physical activity levels and weaker daily rhythmicity. Furthermore, our UTMOST cross-tissue transcriptome-wide association analyses also implicate that the adipose tissues and skeletal muscle tissues, which are mostly related to metabolism and physical activity, may have important roles as they together with the brain cortex are most enriched. Our study suggests the complex interplay among activity, circadian rhythm, metabolic phenotypes and the central nervous system.

We also note that the gene TREH, which was identified in UTMOST tissue enrichment analysis and whose function is related to digestion and galactose metabolism, is associated with sleep start in esophagus, stomach and colon that all belong to the digestive system, suggesting the link between digestion and sleep. From daily experience, a heavy dinner or an empty stomach may affect the ability to fall asleep, and on the other hand, insomnia may also affect the digestive system. It is known from previous studies^68–70^ that sleep problems are associated with gastrointestinal problems and digestive disorders such as gastroesophageal reflux disease and irritable bowel syndrome. We are just beginning to dissect the underlying genetic architecture of sleep, and details on the complex relationships of sleep with the central nervous system and the metabolic system remain to be studied.

This study demonstrates the utility of device measures by presenting applications of population-level objective physical activity data in genetic studies, using novel methods to effectively extract sleep, activity and daily rhythm features. Our results show that accelerometer-derived sleep duration and sleep start/end correlate well with self-reported sleep duration, chronotypes, and hypersomnia. The discrepancy between accelerometer-derived and self-reported physical activity measures are likely due to self-report bias, as people tend to overestimate their daily activity and underestimate sedentary behaviors. The HMM-based algorithm can classify sleep/wake epochs, and the estimated HMM parameters can further be used as sleep and activity related features directly: mean activity levels and activity variability during sleep or wake can characterize individual sleep-activity patterns effectively^71^. In a similar manner, the Penalized Multi-band Learning approach^39^ can identify the population-level dominant periodic information that depicts daily rhythms, and further, individual variations in the strength of periodic signals can be utilized as circadian and daily rhythm features in further analyses. Our study is the first to extract and utilize periodic features from objective physical activity data in genetic studies to examine the genetic architecture of circadian and daily rhythms.

Our study promotes the utility of objective physical activity measures in sleep and physical activity studies when coupled with automated algorithms. The HMM-based sleep/wake identification algorithm and the Penalized Multi-band Learning approach are particularly useful in large-scale population studies when additional sleep validation data are unavailable and manual data examination is labor-intensive and not feasible. More statistical methods are in need to expand and promote the application of actigraphy in clinical and epidemiological studies. Our current circadian study focuses on periodicities, and we will develop new methodology such as functional analysis to extract new dimensions of information and fully exploit actigraphy data in our future work.

In summary, using large-scale population studies like the UK Biobank study with objective physical activity data and genetic data available, we were able to extract meaningful sleep, activity and circadian variables from time-series data using automated algorithms and further conduct genetic analysis to deepen our understanding of the underlying genetic structure. Our study demonstrates the effectiveness of our methods and the utility of device-based activity data in sleep and circadian studies. Our methods can help expand the application of wearable device data in health studies and further provide novel insights into the shared genetic architectures of sleep, activity, and circadian rhythms with metabolic and neurological traits.

## Methods

### Data

Data were collected from the UK Biobank study^72^, a longitudinal population-based study with around 500,000 participants living in the UK. Genetic data from 487,409 participants were available when we accessed the UK Biobank data^72, 73^ in September, 2017. The dataset includes around 96 million single nucleotide polymorphisms (SNPs), including imputation based on the UK10K haplotype, 1000 Genomes Phase 3, and Haplotype Reference Consortium (HRC) reference panels^73^. We applied filters to exclude SNPs with Minor Allele Frequency < 0.1%, Hardy-Weinberg equilibrium < 1e-10 and imputation quality score (UKBB information score^73^) < 0.8. After these steps, a total of 11,024,754 SNPs remained in further analyses.

Besides genetic data, a subset of 103,712 UK Biobank participants agreed to have their objective physical activity data^74^ collected, and they were asked to wear an Axivity AX3 wrist accelerometer for seven consecutive days from 2013 to 2015. The device has been demonstrated to provide equivalent output to the GENEActiv accelerometer, which has been validated against free-living energy expenditure assessment methods.^75–77^ This study was covered by the general ethical approval for UK Biobank studies from the National Research Ethics Service by National Health Service on 17th June 2011 (Reference 11/NW/0382). The accelerometer dataset that we acquired in September of 2017 consists of data in the activity count format summarized every five-second epoch from 103,706 participants. We applied similar quality control procedures as other accelerometer studies^78^. We excluded individuals with flagged data problems, poor wear time, poor calibration, recorded interrupted periods, or inability to calibrate activity data on the device worn itself requiring the use of other data. We also excluded individuals if the number of data recording errors was greater than 3rd quartile + 1.5×IQR. After pre-processing, 92,631 individuals remained, and 90,515 of them had genotyping data that were available for further genetic analyses.

### Identification of Loci Associated with Sleep and Circadian Rhythms

Sleep and circadian rhythm phenotypes were derived from accelerometer data. For sleep, because it is a large-scale population study and no validation sleep logs are available, we need unsupervised algorithms to extract sleep features from accelerometer data. Specifically, we developed an unsupervised sleep-wake identification algorithm^71^ based on Hidden Markov Model (HMM) ^79, 80^ to infer the sequence of “hidden states” of sleep or wake for each individual. This HMM can be directly applied to summary activity count data, which are widely used activity data formats that can save storage space, lengthen the duration of use of wearable devices, and increase computational efficiency. HMM assumes that in the sleep state or the wake state, activity counts follow different distributions as activity counts tend to be fewer in the sleep state compared to those in the wake state^71^. We used log transformation of activity counts and assumed that they follow different Gaussian distributions to infer the sequence of “hidden states” ^71^.

In our model, we consider the log transformed data: log(count+1) as the observed data, because the large range of observed activity counts from zero to several thousand per epoch poses both statistical and computational challenges in data analysis. Our empirical results suggest that the HMM algorithm works well for the log transformed data. We observe activity count data from time 1 to time *T: O*^(*T*)^ = {*O*_1_, *O*_2_, …, *O_T_*}. Let *X*^(*T*)^ = {*X*_1_, *X*_2_,…, *X_T_*} denote the sequence of the corresponding hidden states across these T time points, where each X_i_ can be one of the two possible hidden states *S* = {*s*_1_,*s*_2_} in each epoch, with *s*_1_ denoting the sleep state and *s*_2_ denoting the wake state. We assume that X_i_ follows a Markov model, that is the hidden state *X*_*t*+1_ at time *t*+1 solely depends on *X_t_*, and the observation *O_t_* at time *t* solely depends on the hidden state *X_t_*.

*A* denotes the 2 by 2 transition probability matrix, in which *a_ij_* represents the transition probability from state *s_i_* to state *S_j_*. The emission probability *P*(*O_t_*| *X_t_*) denoted by *B* depends on the state of *X_t_*. If *X_t_* = *s*_1_ in the sleep state, we assume that the log transformed count follows zero-inflated truncated Gaussian distribution, which is truncated from 0 to the left. It has a zero component because sleep is associated with rare movements and activity measurements during sleep often involve many zeros. Therefore:

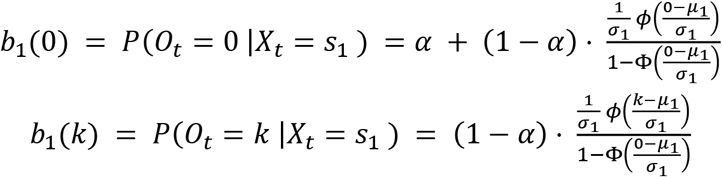

where *α* is the probability of extra zeros, is the mean, *σ*_1_ is the standard deviation, *φ*(·) is the probability density function of the standard normal distribution, and Φ(·) is its cumulative distribution function.

If *X_t_* = *s*_2_ in the wake state, we assume that the log transformed count follows the Gaussian distribution:

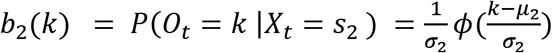

where *μ*_2_ is the mean and *σ*_2_ is the standard deviation of the Gaussian distribution.

Therefore, the set of parameters for the emission probability is *B* = {*B*_1_, *B*_2_} = {*α, μ*_1_, *σ*_1_, *μ*_2_, *σ*_2_}. To initiate the Markov chain, we also need the initial state probabilities Π = {*π*_0_, *π*_1_}. HMM can be fully specified by Θ = {*A,B*, Π}. To obtain Θ* = argmax_Θ_ P{ *O*^(*T*)^| Θ }, we can use the Baum-Welch algorithm, and we further look for the optimal path of hidden states *X*^(*T*)*^ = argmax_*X*_^(*T*)^ *P*{*X*^(*T*)^, *O*^(*T*)^| Θ* } using the Viterbi algorithm^81^. The optimal hidden states *X*^(*T*)*^ are exactly the sequence of inferred sleep/wake states.

Besides the inferred sequence of sleep/wake states, the HMM parameters estimated for each individual, including mean activity levels and variability (standard deviation) for sleep and wake states, can characterize individual sleep and activity behaviors and therefore were used as sleep and activity phenotypes. In addition, we inferred sleep duration, sleep start and end based on the inferred sequence of sleep/wake states. We created two categorical variables as sleep duration phenotypes indicating whether the sleep duration is < 5 hours or > 10 hours, the thresholds of which come from the National Sleep Foundation^82^ and are not recommended for middle-age and older adults. The inferred sleep start time and sleep end time are used as phenotypes for the timing of sleep onset and wake-up. Sleep start and sleep end are related to but not exactly the same as chronotypes, which describe whether a person is a morning person, getting up early and remaining more active in the day, or a night person, remaining more active later of the day and staying up late at night, and chronotypes are usually measured in self-reported questionnaires. Published work^83^ using UK Biobank data has utilized midpoint of sleep, the least active 5 hours of the day, as sleep timing to be compared with self-reported chronotypes. Here we did not replicate the study but examined timing of sleep onset and wake-up, as for different types of sleep disorders some people have difficulty falling asleep while others find it problematic to wake up too early^84^.

For circadian rhythm characteristics, we derived circadian features by utilizing the Penalized Multi-Band Learning (PML) approach^39^, which extracts periodic information using Fast Fourier Transform (FFT) and then performs penalized selection based on regularization, a classic approach used in machine learning^85, 86^, to identify population-level dominant periodicities such as 1-day, 1/2-day, and 1/3-day periodicities that can characterize daily activity rhythms. The strengths of FFT signals at dominant periodicities are then used as circadian phenotypes in genetic analysis.

The PML algorithm is as follows. Let matrix *X* ∈ *R^n × p^*, where *n* denotes the number of individual observations, and *p* denotes the number of periodicities from FFT. Specifically, *X* = (*x*_1_, *x*_2_, …, *x_p_*), where *X_j_* is the vector of length *n* for the *j*th periodicity.

Let Θ be the diagonal matrix selecting columns from *X* such that 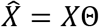:

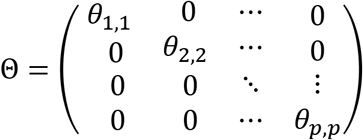

where 0 ≤*θ_ij_*,≤1, *j* =1,…, *p*. Θ identifies columns of dominant periodicities from *X* in the way that dominant periodicities corresponding to nonzero *θ_j,j_*’s are selected. We minimize the squared Frobenius norm 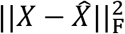, and by using properties of the Frobenius norm, we can get:

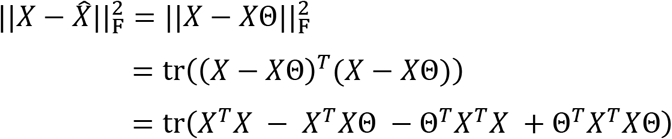

Because is fixed, it is equivalent to minimize:

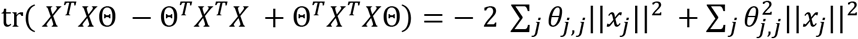

In order to estimate Θ and identify dominant periodicities, we use a penalized selection method similar to Lasso, a widely used method in shrinkage and feature selection in regression models that is most effective in selecting a few important features while suppressing other non-selected features to 0^86^. In our case, Lasso penalty serves to select a few dominant periodicities through diagonal elements of Θ instead of regression coefficients. Further, we add an elastic-net like penalty term onto the Frobenius norm, namely a combination of L1 and L2 norms^85^:

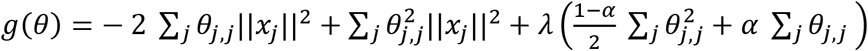

where *λ* is the tuning parameter and *α* controls the balance between the L1 and L2 norms. Note that *θ_j,j_*’s are nonnegative and thus we do not need to take the absolute value for the L1 norm. By setting *λ* large enough, all diagonal elements of Θ, namely all *θ_j,j_*’s, are suppressed to zero and no periodicities are selected. As *λ* decreases, some *θ_j,j_*’s become nonzero and they correspond to the most dominant periodicities that are selected sequentially according to how dominant they are.

To minimize *g*(0), we take the partial derivative of *g*(0) with respect to each 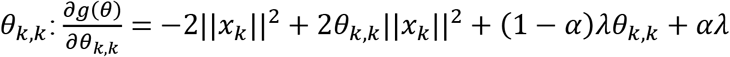 which is convex and also subject to the constraint 0 ≤ *θ_k,k_* ≤ 1. Thus, we have:

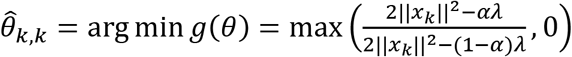

If we only have the L1 penalty *α* = 1 and 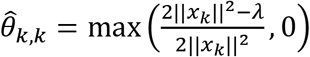. In our case, we use Lasso L1 penalty alone and train *λ,* because we want to select the most important periodicities while suppressing other periodicities to 0.

We use mean squared error (MSE), which is equivalent to the squared Frobenius norm 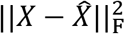, as the criterion for choosing *λ* and the number of nonzero *θ_j,j_*’*s* (the number of dominant periodicities selected). We train *λ* from 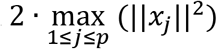 to 0, as 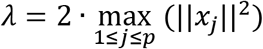 suppresses all *θ_j,j_*’s to 0 and *λ* = 0 gives no penalty. By decreasing *λ*, we identify dominant periodicities sequentially to characterize the daily sleep-activity rhythm. An R package named PML has been developed (https://CRAN.R-project.org/package=PML) for the implementation of the PML algorithm^87^.

To identify genetic loci associated with each sleep and circadian phenotype, we conducted genome-wide association analysis, using PLINK (version 1.9) ^88^. We included age, sex, and the first 20 principal components as covariates when fitting linear models, and we report statistically significant loci using a traditional threshold of 5×10^-8^. We also highlight results from a more stringent threshold of 5×10^-9^ to take into account multiple testing using Bonferroni correction, which is the same threshold suggested in previous studies^67, 89^.

### Genetic Architecture of Sleep and Circadian Rhythms

To estimate heritability^14^, we applied LD score regression^14, 49^ with the LDSC tool. We also conducted partitioned heritability analysis^49^ across tissue categories using the same tool. Significant enrichments for individual traits were identified using the Bonferroni corrected threshold of *p* < 5 × 10^-4^ (10 traits ×10 tissue types).

### Genetic Correlation of Sleep and Daily Rhythms with Other Traits

To examine the genetic correlation of sleep-activity traits and circadian rhythm traits with other traits and diseases, we downloaded GWAS summary statistics in the second round of results from the Neale lab released on August 1, 2018 (http://www.nealelab.is/uk-biobank/). Specifically, we chose activity related traits including time spent doing moderate or vigorous physical activity, screen exposure traits including time spent watching television, computer, or mobile phone, sleep traits such as self-reported sleep duration, chronotypes, and insomnia, mental health traits related to anxiety and depression, BMI and diet traits, alcohol consumption traits, shift-work traits, and respiratory disease treats. The detailed list of 53 traits are shown in Supplementary Table S3. We calculated cross-trait genetic correlation using GNOVA^90^ and highlight the trait-pairs with Bonferroni corrected p-values < 7.9 × 10^-5^ (12 sleep and circadian traits and 53 other traits).

For significantly correlated trait-pairs, we further investigated causal relationships by conducting bi-directional Mendelian Randomization (MR) analyses. We did not analyze trait-pairs related to sleep and activity, which have been studied elsewhere^52, 67^, but primarily focused on the circadian rhythm. We performed two-sample MR analysis using publicly available GWAS summary data extracted from the MR-Base web platform^91^. We used leave-one-out analysis and single-SNP analysis as sensitivity analyses and considered the inverse-variance weighted (IVW) approach, MR-Egger^92^, weighted median estimation and weighted mode estimation methods to examine whether there are consistent MR results across methods.

### Cross-Tissue Transcriptome-Wide Association Analysis

To investigate functional and biological mechanisms underlying sleep-activity and circadian rhythms, we applied UTMOST^53^ that utilizes GWAS summary statistics and integrates eQTL information to perform tissue enrichment analysis in 44 tissue types and identify single-tissue and cross-tissue gene-trait associations. The cross-tissue gene-trait association is evaluated via a joint test summarizing single-tissue association statistics and quantifying the overall gene-trait association. The UTMOST^53^ p-value threshold after Bonferroni correction for 15,120 genes is 3.3× 10^-6^. Gene ontology enrichment analysis was further conducted using DAVID^61^.

## A Supplementary Materials

### A.1 Supplementary Figures

**Figure S1.**
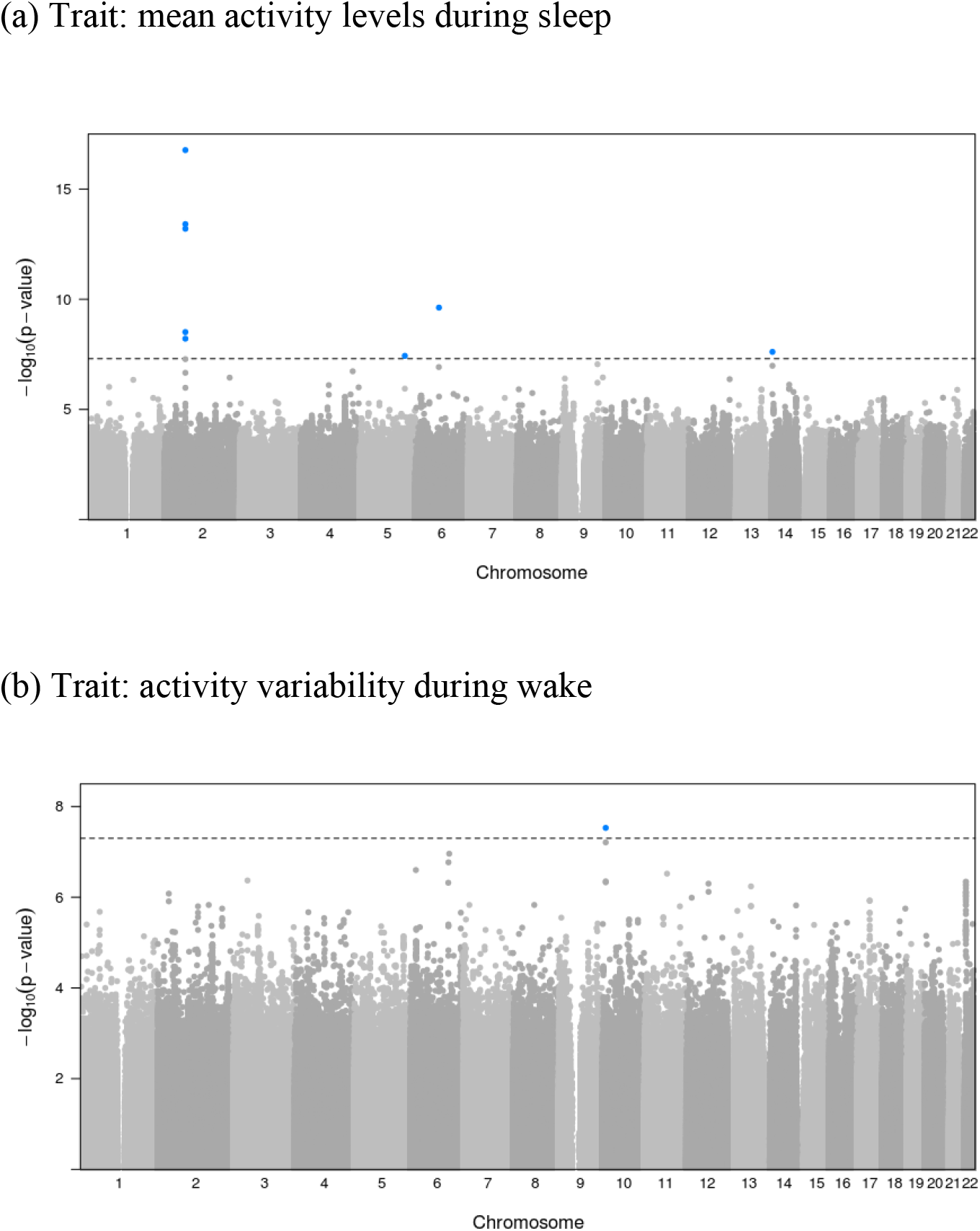
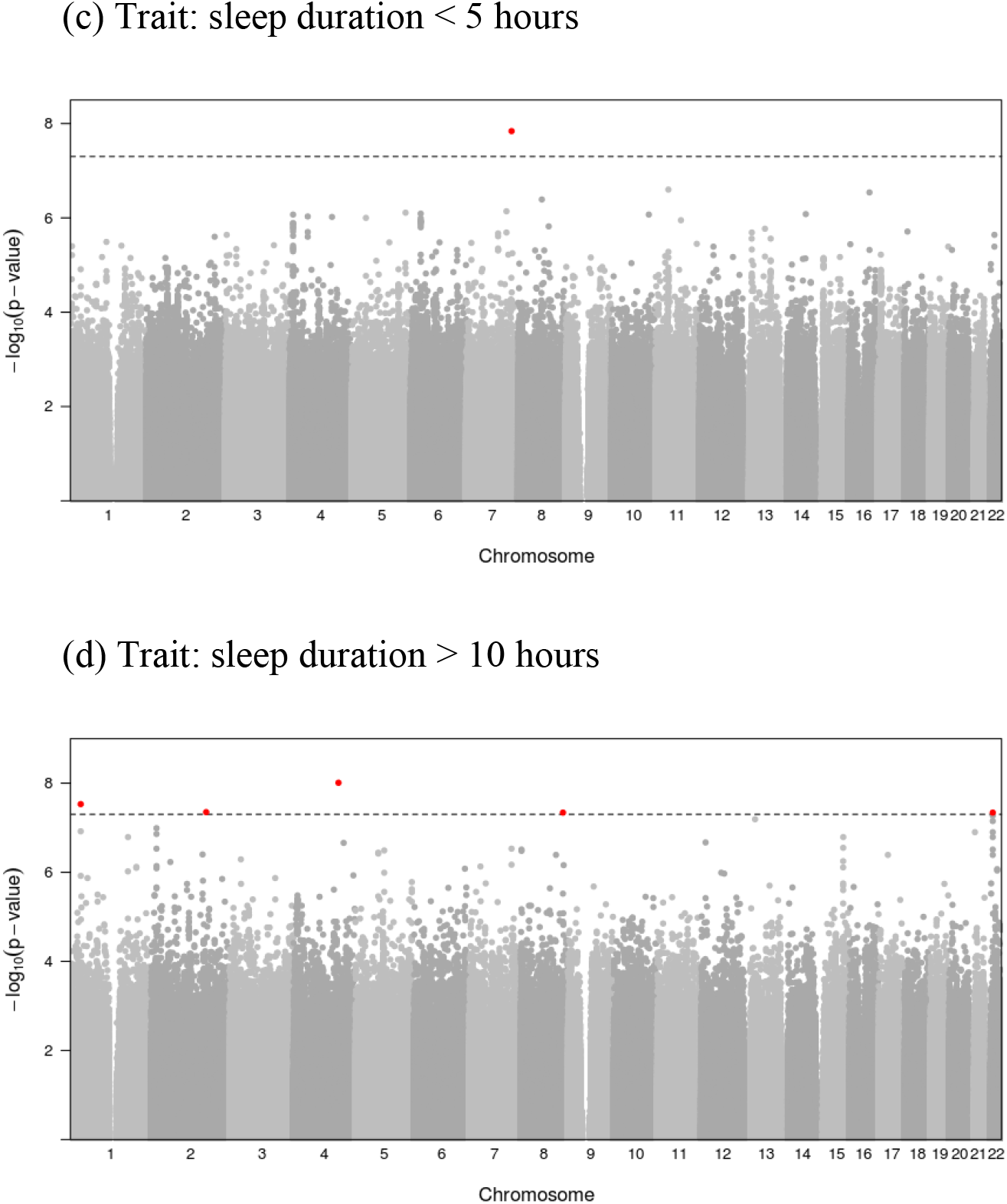
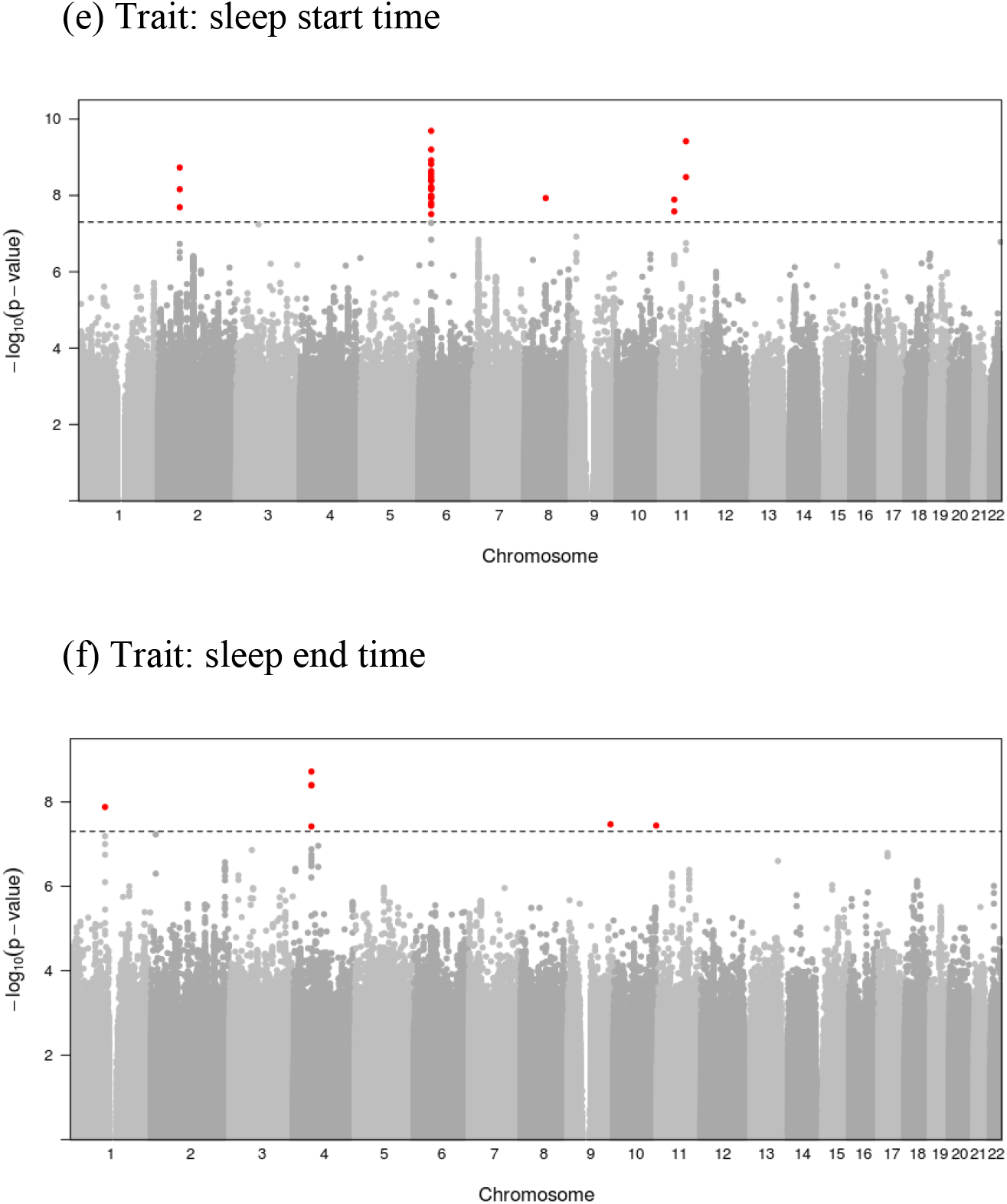
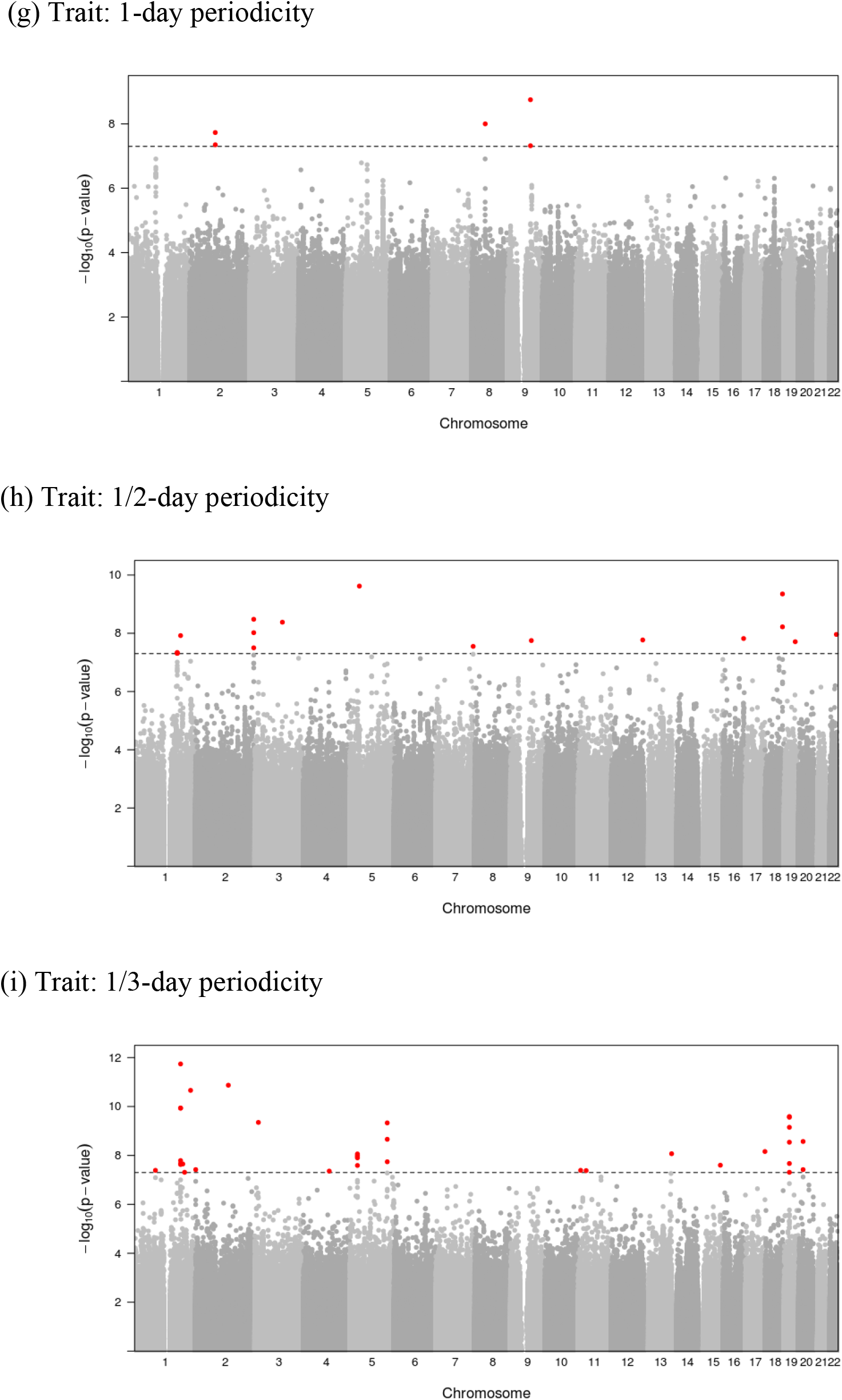
Manhattan plots for genome-wide association studies of sleep and activity traits, derived sleep traits, and circadian traits.

**Figure S2.**
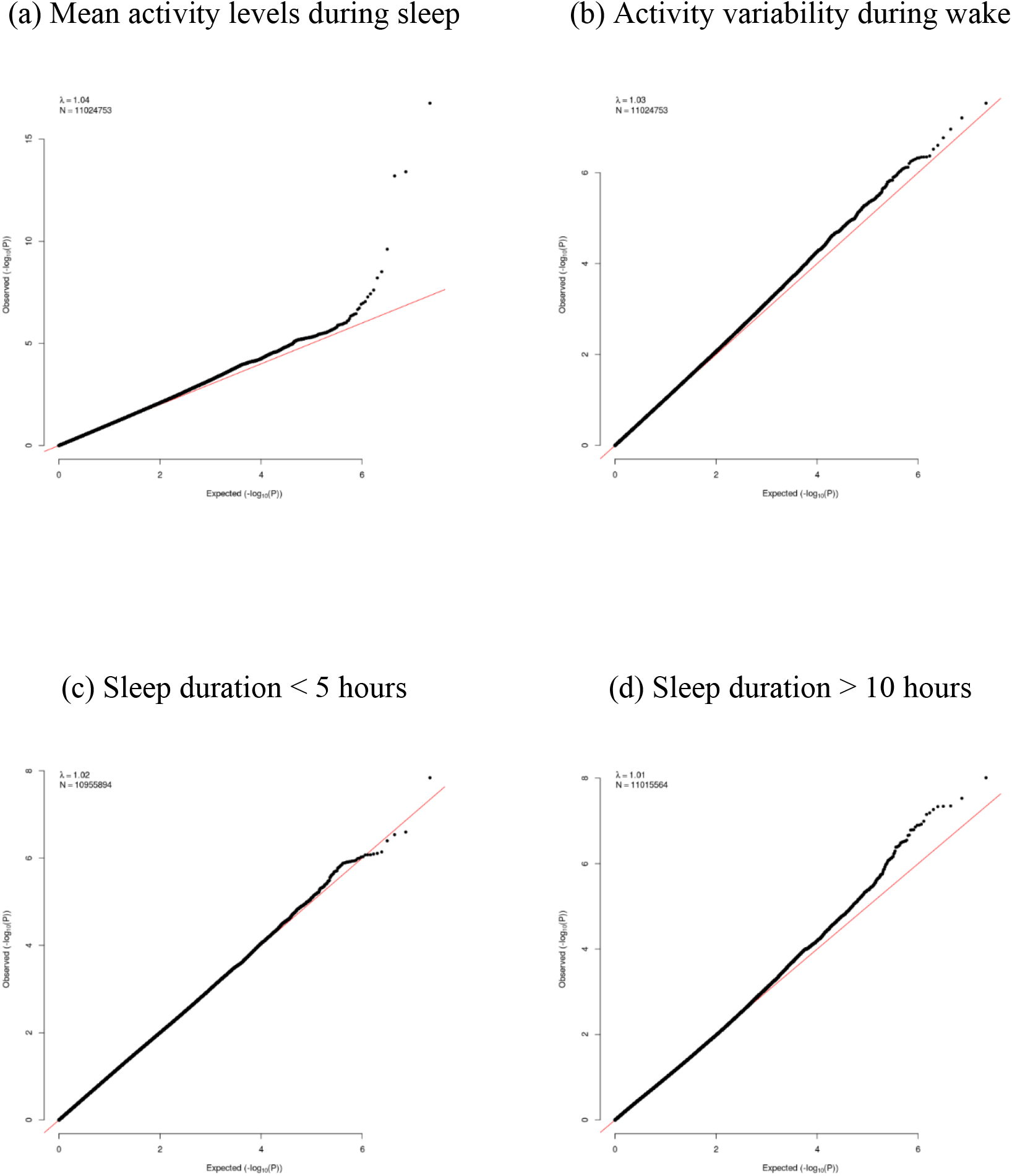
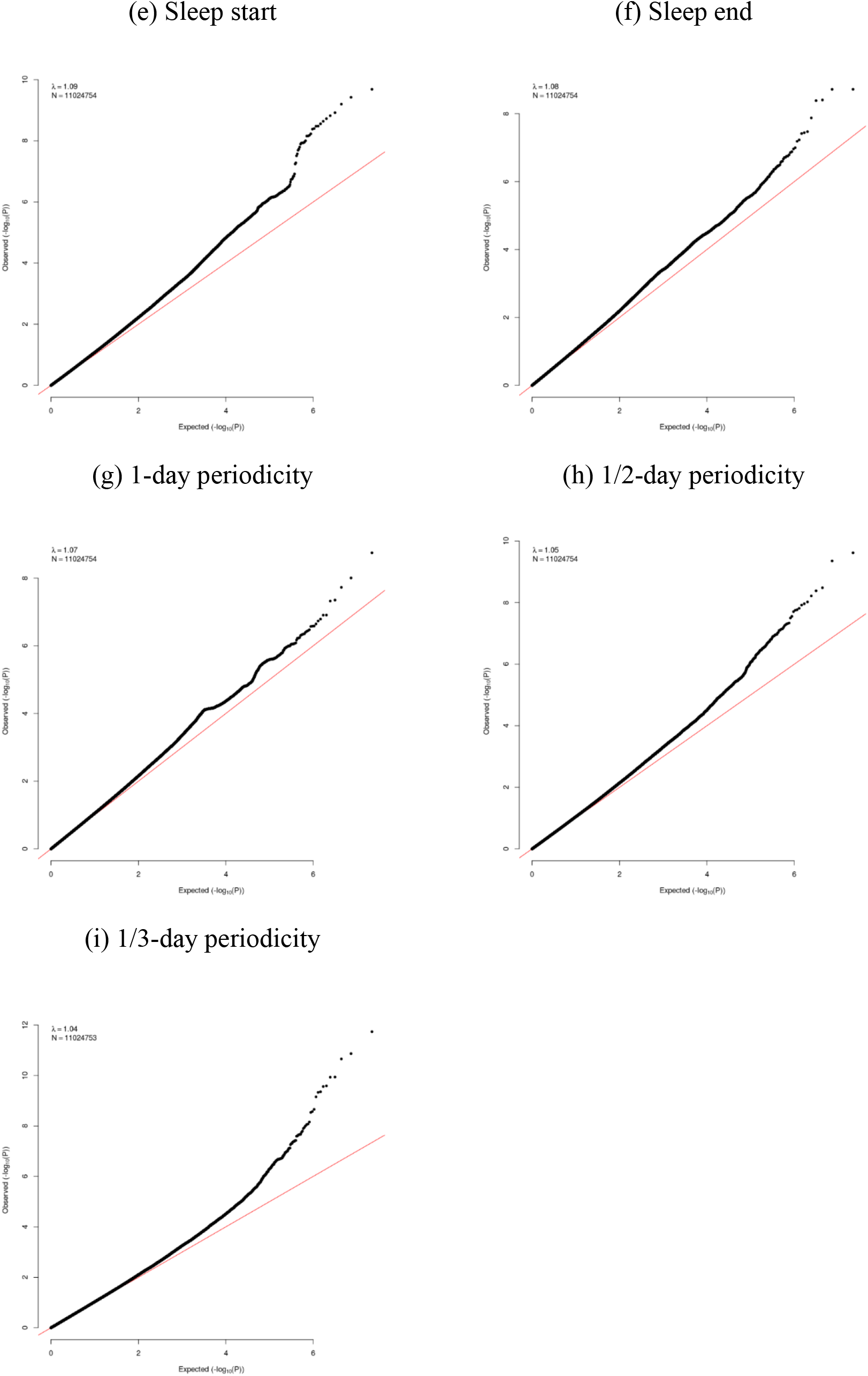
QQ-plots for checking population stratification in genome-wide association studies.

**Figure S3.**
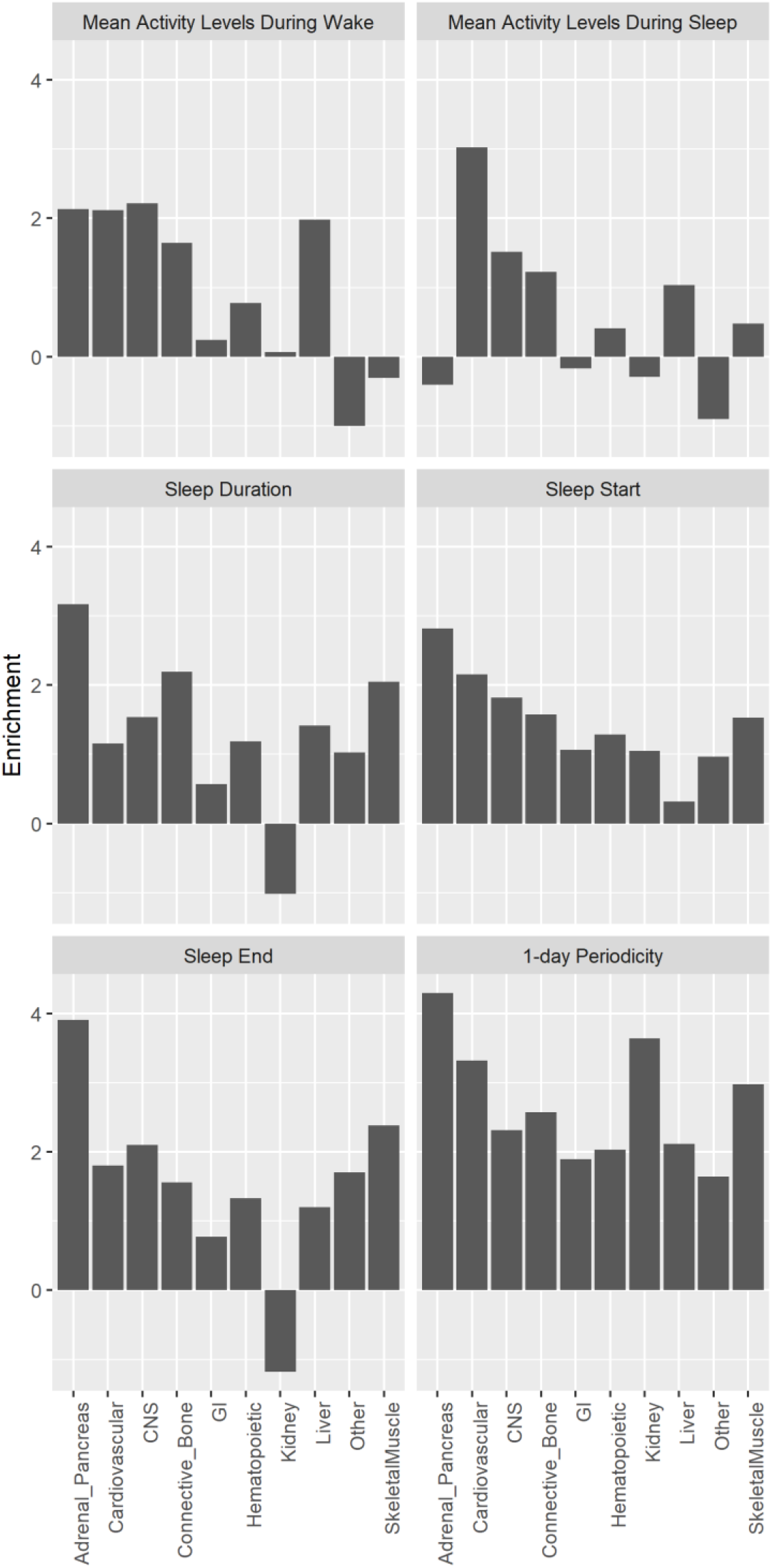
Partitioned heritability enrichment analysis across 10 broad tissue types for activity, sleep time, and circadian rhythm traits among 90,515 UK Biobank participants with objective physical activity measurements.

**Figure S4.**
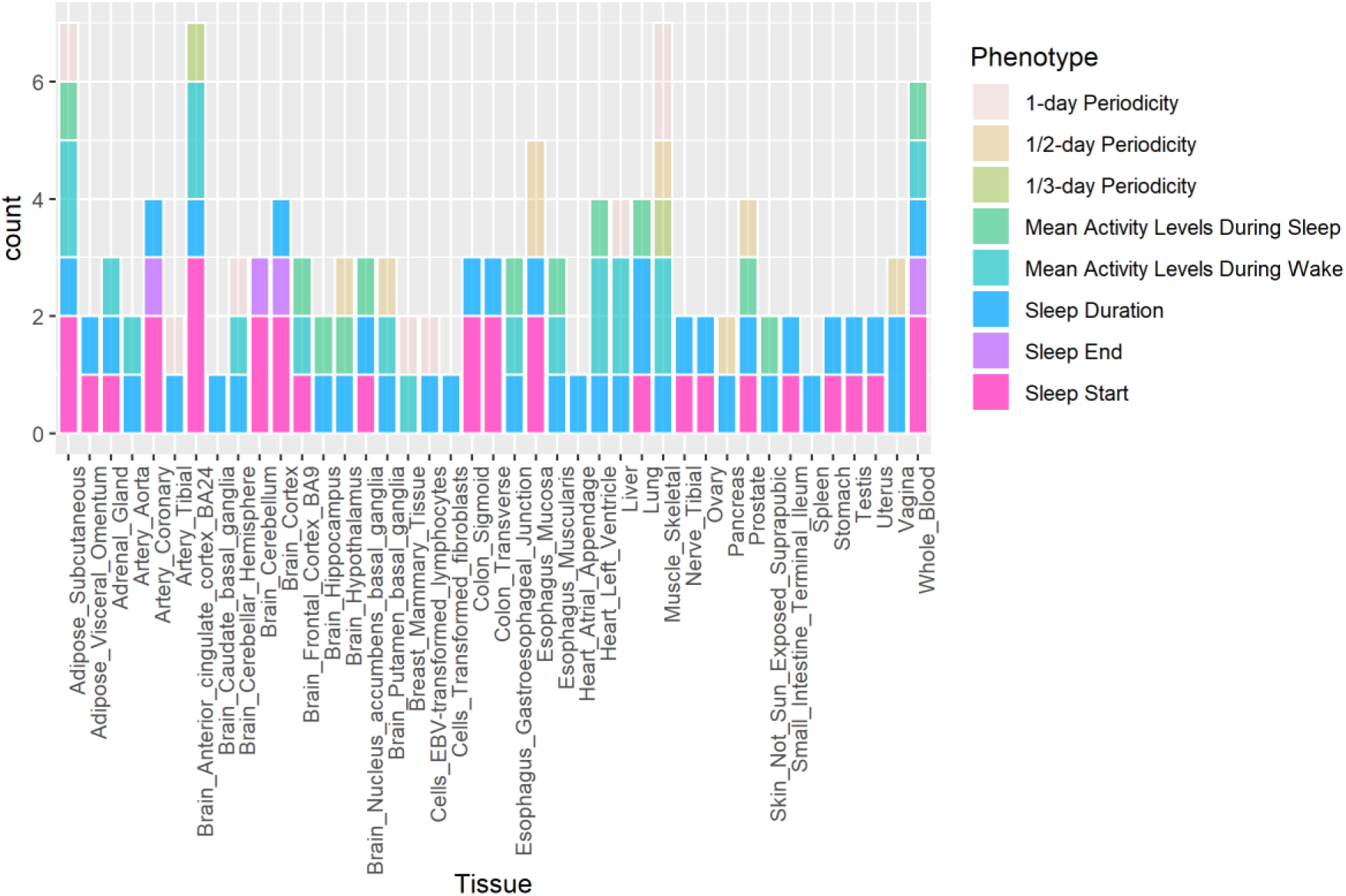
Tissue enrichment analysis for gene-trait pairs across 44 tissue types for activity, sleep time, and circadian rhythm traits extracted from objective physical activity measurements among 90,515 UK Biobank participants.

### A.2 Supplementary Tables

**Table S1.** The SNPs identified in genome-wide association studies at the significance level of 5 × 10^-8^ that are associated with sleep start and sleep end traits inferred from accelerometer-measured physical activity in 90,515 UK Biobank participants.

| Trait | Chr | Position | ID | Function | Nearest Gene | Risk Allele | BETA | SE | P |
| --- | --- | --- | --- | --- | --- | --- | --- | --- | --- |
| Sleep Start | 2 | 66750564 | rs113851554 | intronic | MEIS1 | T | 0.070 | 0.012 | 1.88E-09 |
|  | 2 | 66785180 | rs11679120 | intronic | MEIS1 | A | 0.075 | 0.013 | 6.97E-09 |
|  | 2 | 66799986 | rs11693221 | downstream | MEIS1(dist=95) | T | 0.072 | 0.013 | 2.04E-08 |
|  | 6 | 38399009 | rs9349081 | intronic | BTBD9 | A | -0.036 | 0.006 | 1.16E-08 |
|  | 6 | 38401926 | rs78054489 | intronic | BTBD9 | T | -0.036 | 0.006 | 1.10E-08 |
|  | 6 | 38402197 | rs62397049 | intronic | BTBD9 | G | -0.035 | 0.006 | 1.17E-08 |
|  | 6 | 38404140 | rs4714160 | intronic | BTBD9 | T | -0.037 | 0.006 | 3.97E-09 |
|  | 6 | 38404333 | rs9349082 | intronic | BTBD9 | A | -0.037 | 0.006 | 3.32E-09 |
|  | 6 | 38405652 | rs62397050 | intronic | BTBD9 | C | -0.037 | 0.006 | 4.14E-09 |
|  | 6 | 38416590 | rs9380753 | intronic | BTBD9 | A | -0.037 | 0.006 | 2.29E-09 |
|  | 6 | 38418303 | rs4714162 | intronic | BTBD9 | C | -0.037 | 0.006 | 2.77E-09 |
|  | 6 | 38432656 | rs62397055 | intronic | BTBD9 | T | -0.037 | 0.006 | 9.94E-09 |
|  | 6 | 38437303 | rs9369062 | intronic | BTBD9 | C | -0.035 | 0.006 | 6.29E-10 |
|  | 6 | 38438771 | rs4714163 | intronic | BTBD9 | C | -0.035 | 0.006 | 1.51E-09 |
|  | 6 | 38440970 | rs3923809 | intronic | BTBD9 | G | -0.033 | 0.006 | 6.95E-09 |
|  | 6 | 38444040 | rs6920488 | intronic | BTBD9 | G | -0.035 | 0.006 | 1.20E-09 |
|  | 6 | 38447870 | rs13219518 | intronic | BTBD9 | T | -0.037 | 0.006 | 2.06E-10 |
|  | 6 | 38454433 | rs9349087 | intronic | BTBD9 | G | -0.034 | 0.006 | 5.90E-09 |
| Sleep Start | 6 | 38466562 | rs9349088 | intronic | BTBD9 | G | -0.034 | 0.006 | 6.59E-09 |
|  | 6 | 38469187 | rs10947738 | intronic | BTBD9 | C | -0.033 | 0.006 | 1.86E-08 |
|  | 6 | 38469323 | rs10947739 | intronic | BTBD9 | T | -0.033 | 0.006 | 1.60E-08 |
|  | 6 | 38470087 | rs4236060 | intronic | BTBD9 | T | -0.033 | 0.006 | 3.09E-08 |
|  | 8 | 65505208 | rs77576509 | UTR3 | CYP7B1(NM_004820:c.*3991G>C) | C | -0.496 | 0.087 | 1.16E-08 |
|  | 11 | 43671680 | rs79899879 | intergenic | MIR129-2(dist=68647),HSD17B12(dist=30463) | C | 0.470 | 0.084 | 2.65E-08 |
|  | 11 | 43673370 | rs77677460 | intergenic | MIR129-2(dist=70337),HSD17B12(dist=28773) | G | 0.471 | 0.083 | 1.27E-08 |
|  | 11 | 80275391 | rs76322105 | intergenic | LOC101928944(dist=186853) | C | 0.528 | 0.084 | 3.76E-10 |
|  | 11 | 80275893 | rs7109066 | intergenic | LOC101928944(dist=186351) | G | 0.478 | 0.081 | 3.29E-09 |
| Sleep End | 1 | 107738826 | rs60650667 | intronic | NTNG1 | C | 0.512 | 0.090 | 1.33E-08 |
|  | 4 | 55435875 | rs17084481 | intergenic | LINC02260(dist=33503) | T | -0.451 | 0.082 | 3.83E-08 |
|  | 4 | 55442330 | rs17084486 | intergenic | LINC02260(dist=27048) | A | -0.502 | 0.085 | 3.94E-09 |
|  | 4 | 55445652 | rs6833573 | intergenic | LINC02260(dist=23726) | C | -0.515 | 0.086 | 1.91E-09 |
|  | 4 | 55447888 | rs6817676 | intergenic | LINC02260(dist=21490) | T | -0.513 | 0.085 | 1.91E-09 |
|  | 4 | 55467932 | rs114260939 | intergenic | LINC02260(dist=1446) | T | -0.525 | 0.089 | 4.11E-09 |
|  | 9 | 132183853 | rs28361713 | intergenic | LINC00963(dist=67086) | A | -0.508 | 0.092 | 3.37E-08 |
|  | 10 | 132361598 | rs74162666 | intergenic | GLRX3(dist=382952) | A | 0.499 | 0.091 | 3.63E-08 |

**Table S2.**
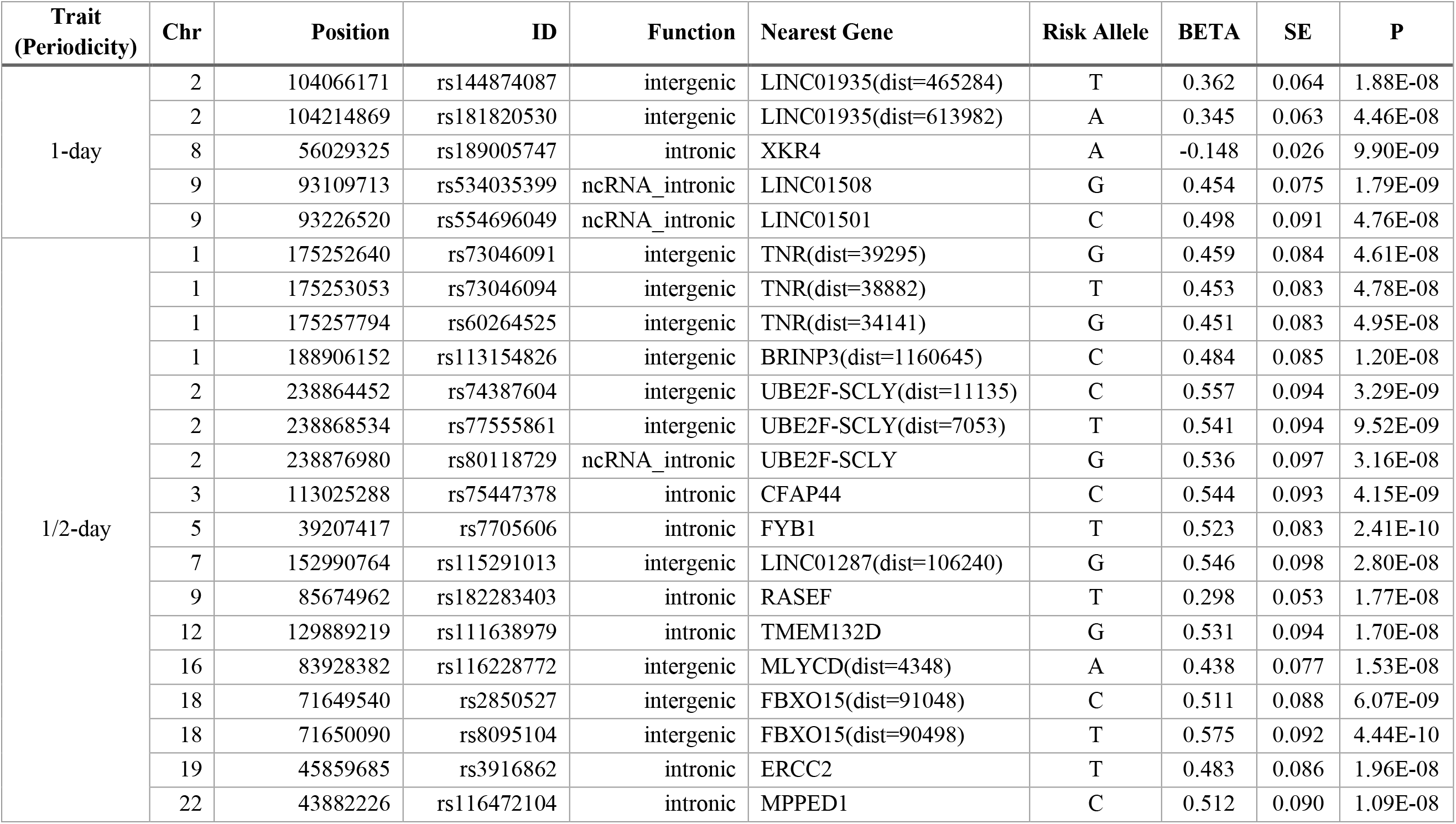
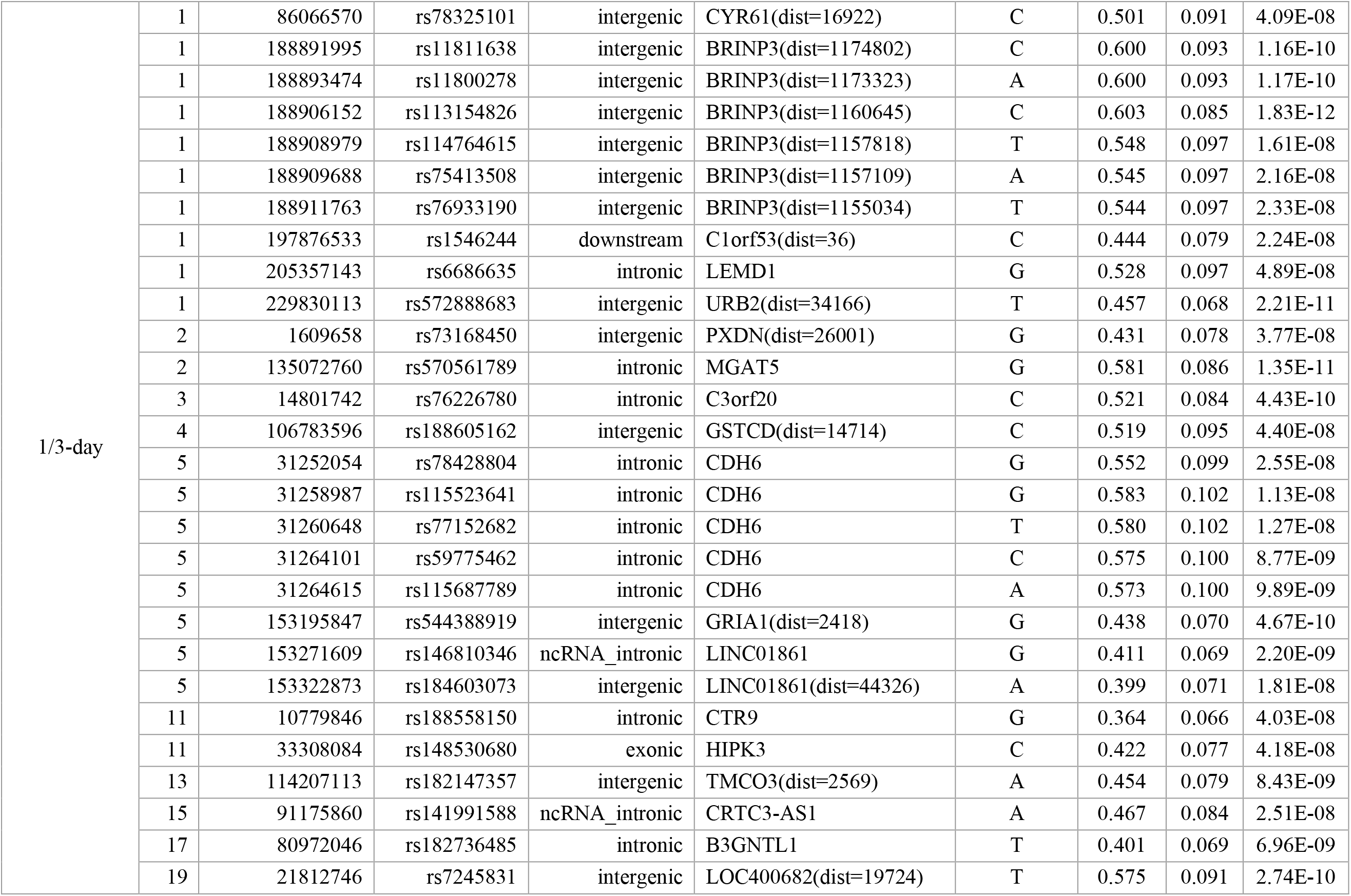
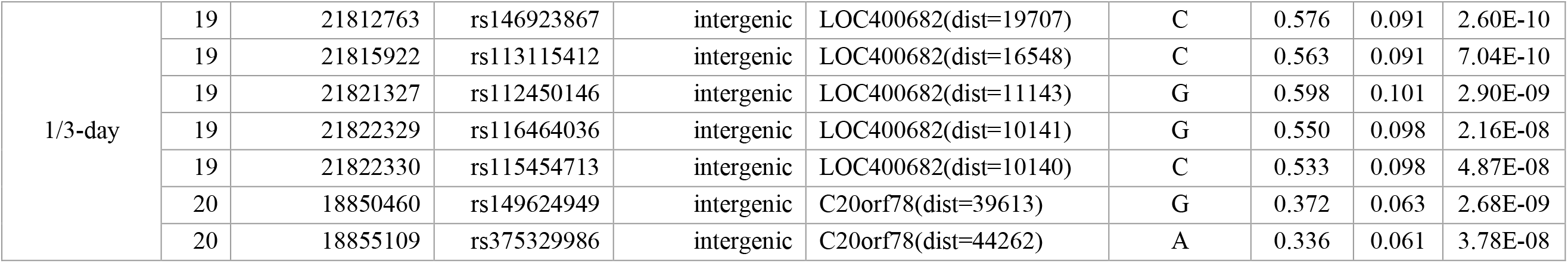
The SNPs identified in genome-wide association studies at the significance level of 5 × 10^-8^ that are associated with dominant periodicities as circadian traits inferred from accelerometer-measured physical activity in 90,515 UK Biobank participants.

**Table S3.**
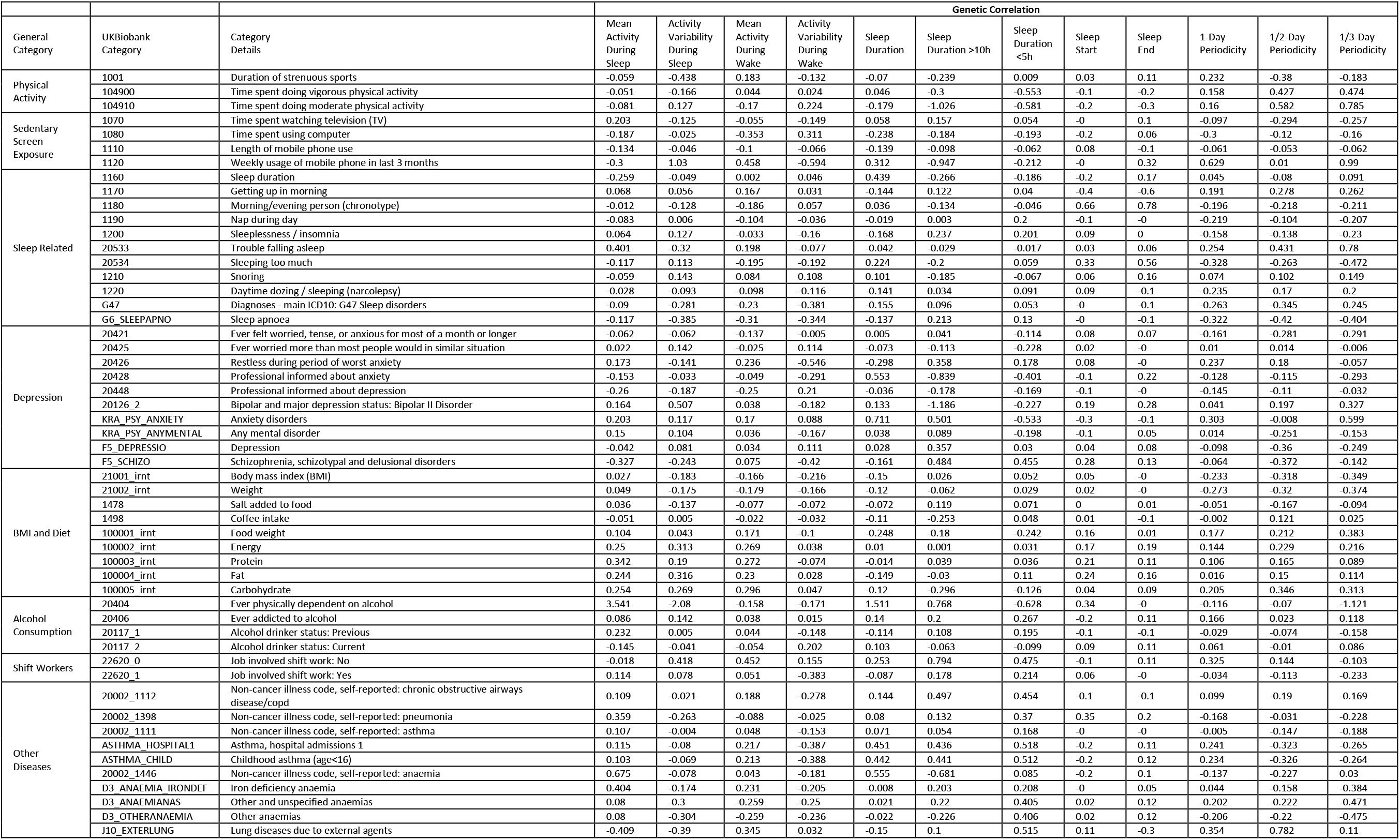
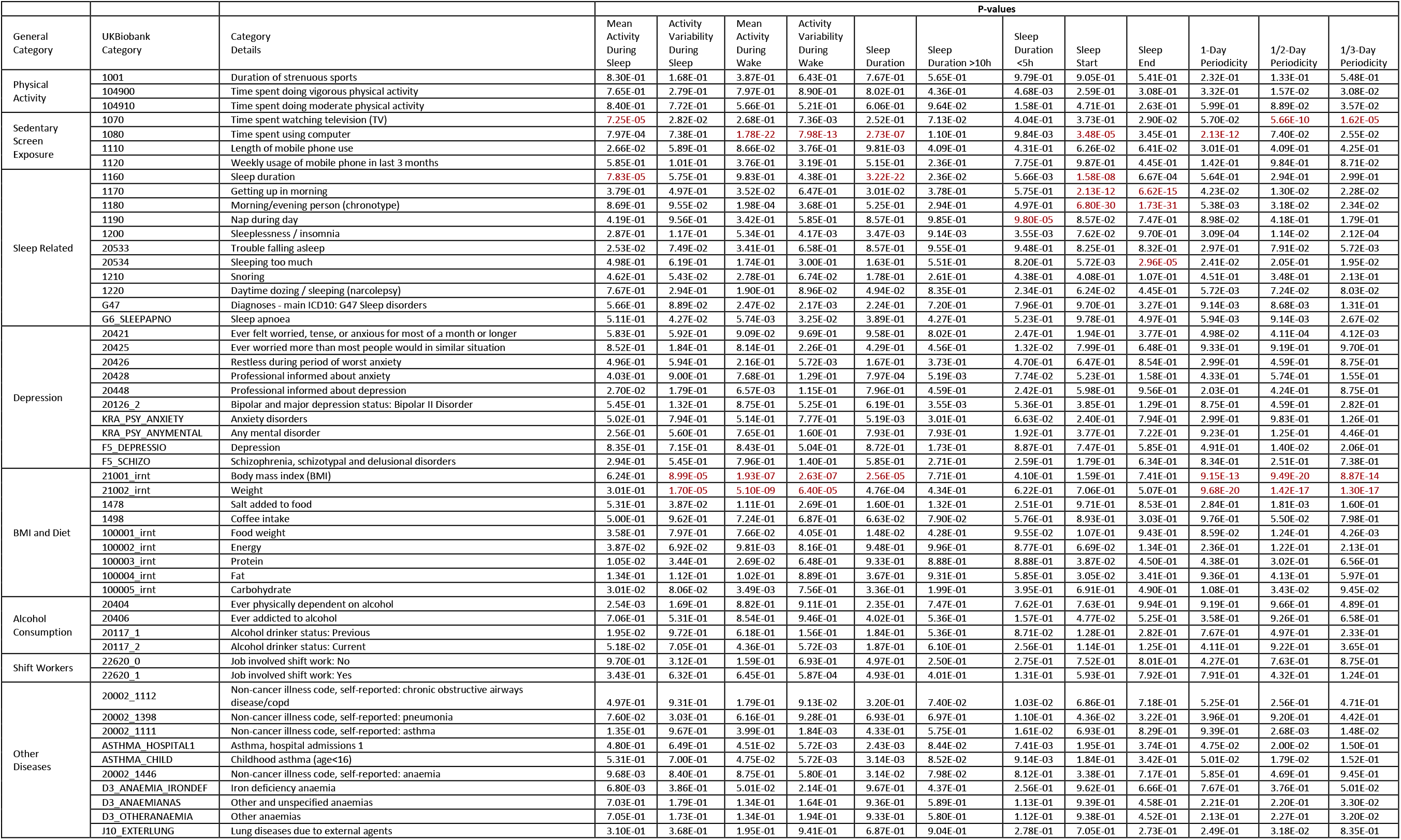
Estimated genetic correlation of sleep and circadian traits with other complex traits and the corresponding p-values.

**Table S4.** Single-tissue gene-trait association test results from tissue enrichment analysis using UTMOST.

| Phenotype | Tissue | GENE | zscore | pvalue |
| --- | --- | --- | --- | --- |
| Mean Activity Levels During Sleep | Esophagus_Muscularis | BRSK1 | 4.63 | 3.70E-06 |
| Mean Activity Levels During Sleep | Prostate | BRSK1 | 4.80 | 1.57E-06 |
| Mean Activity Levels During Sleep | Skin_Not_Sun_Exposed_Suprapubic | BRSK1 | 4.71 | 2.52E-06 |
| Mean Activity Levels During Sleep | Brain_Frontal_Cortex_BA9 | CEP70 | -4.59 | 4.46E-06 |
| Mean Activity Levels During Sleep | Esophagus_Gastroesophageal_Junction | CEP70 | -4.68 | 2.84E-06 |
| Mean Activity Levels During Sleep | Brain_Hypothalamus | CEP70 | -4.61 | 4.09E-06 |
| Mean Activity Levels During Sleep | Lung | CEP70 | -4.63 | 3.73E-06 |
| Mean Activity Levels During Sleep | Heart_Left_Ventricle | KATNAL2 | 4.64 | 3.41E-06 |
| Mean Activity Levels During Sleep | Brain_Nucleus_accumbens_basal_ganglia | KCNH7 | -5.01 | 5.35E-07 |
| Mean Activity Levels During Sleep | Adipose_Subcutaneous | L3MBTL2 | -4.75 | 2.07E-06 |
| Mean Activity Levels During Sleep | Whole_Blood | L3MBTL2 | -4.72 | 2.38E-06 |
| Mean Activity Levels During Sleep | Brain_Hippocampus | PLA2G4B | -4.84 | 1.31E-06 |
| Mean Activity Levels During Wake | Brain_Putamen_basal_ganglia | ARID3B | 4.81 | 1.54E-06 |
| Mean Activity Levels During Wake | Heart_Left_Ventricle | BEAN1 | 4.70 | 2.57E-06 |
| Mean Activity Levels During Wake | Adipose_Subcutaneous | GIPC2 | -4.72 | 2.32E-06 |
| Mean Activity Levels During Wake | Heart_Left_Ventricle | GIPC2 | 4.83 | 1.35E-06 |
| Mean Activity Levels During Wake | Breast_Mammary_Tissue | GIPC2 | -4.74 | 2.12E-06 |
| Mean Activity Levels During Wake | Adipose_Subcutaneous | L3MBTL2 | -5.39 | 7.05E-08 |
| Mean Activity Levels During Wake | Muscle_Skeletal | L3MBTL2 | -5.62 | 1.92E-08 |
| Mean Activity Levels During Wake | Liver | MBD2 | 4.78 | 1.71E-06 |
| Mean Activity Levels During Wake | Brain_Anterior_cingulate_cortex_BA24 | PRUNE | -4.67 | 3.02E-06 |
| Mean Activity Levels During Wake | Whole_Blood | SEC11C | -4.70 | 2.63E-06 |
| Mean Activity Levels During Wake | Brain_Anterior_cingulate_cortex_BA24 | SOD3 | -4.75 | 2.06E-06 |
| Mean Activity Levels During Wake | Muscle_Skeletal | SPARC | -5.03 | 4.86E-07 |
| Mean Activity Levels During Wake | Adrenal_Gland | TRAF3 | 4.87 | 1.14E-06 |
| Mean Activity Levels During Wake | Artery_Aorta | TRAF3 | 4.87 | 1.12E-06 |
| Mean Activity Levels During Wake | Brain_Cerebellar_Hemisphere | TRAF3 | 4.67 | 3.01E-06 |
| Mean Activity Levels During Wake | Brain_Frontal_Cortex_BA9 | TRAF3 | 5.11 | 3.25E-07 |
| Mean Activity Levels During Wake | Esophagus_Gastroesophageal_Junction | TRAF3 | 4.77 | 1.85E-06 |
| Mean Activity Levels During Wake | Esophagus_Muscularis | TRAF3 | 4.86 | 1.18E-06 |
| Mean Activity Levels During Wake | Liver | TRAF3 | 4.88 | 1.07E-06 |
| Sleep Duration | Vagina | ADAMTS7 | 4.80 | 1.63E-06 |
| Sleep Duration | Adipose_Subcutaneous | GLTP | 4.66 | 3.13E-06 |
| Sleep Duration | Adipose_Visceral_Omentum | GLTP | -4.76 | 1.90E-06 |
| Sleep Duration | Adrenal_Gland | GLTP | -4.71 | 2.42E-06 |
| Sleep Duration | Artery_Aorta | GLTP | 4.76 | 1.97E-06 |
| Sleep Duration | Artery_Coronary | GLTP | 4.74 | 2.11E-06 |
| Sleep Duration | Artery_Tibial | GLTP | -4.74 | 2.12E-06 |
| Sleep Duration | Brain_Anterior_cingulate_cortex_BA24 | GLTP | -4.75 | 2.07E-06 |
| Sleep Duration | Brain_Caudate_basal_ganglia | GLTP | -4.73 | 2.25E-06 |
| Sleep Duration | Brain_Cerebellar_Hemisphere | GLTP | -4.76 | 1.95E-06 |
| Sleep Duration | Brain_Cortex | GLTP | -4.75 | 2.01E-06 |
| Sleep Duration | Brain_Hippocampus | GLTP | -4.75 | 1.99E-06 |
| Sleep Duration | Brain_Hypothalamus | GLTP | -4.76 | 1.92E-06 |
| Sleep Duration | Brain_Nucleus_accumbens_basal_ganglia | GLTP | -4.70 | 2.55E-06 |
| Sleep Duration | Brain_Putamen_basal_ganglia | GLTP | -4.69 | 2.77E-06 |
| Sleep Duration | Cells_EBV-transformed_lymphocytes | GLTP | 4.61 | 3.97E-06 |
| Sleep Duration | Cells_Transformed_fibroblasts | GLTP | 4.75 | 1.99E-06 |
| Sleep Duration | Colon_Sigmoid | GLTP | 4.73 | 2.26E-06 |
| Sleep Duration | Colon_Transverse | GLTP | -4.76 | 1.94E-06 |
| Sleep Duration | Esophagus_Gastroesophageal_Junction | GLTP | -4.75 | 2.00E-06 |
| Sleep Duration | Esophagus_Mucosa | GLTP | 4.73 | 2.29E-06 |
| Sleep Duration | Esophagus_Muscularis | GLTP | 4.65 | 3.28E-06 |
| Sleep Duration | Heart_Atrial_Appendage | GLTP | 4.63 | 3.68E-06 |
| Sleep Duration | Heart_Left_Ventricle | GLTP | -4.74 | 2.15E-06 |
| Sleep Duration | Liver | GLTP | 4.73 | 2.24E-06 |
| Sleep Duration | Lung | GLTP | -4.75 | 2.01E-06 |
| Sleep Duration | Vagina | GLTP | -4.75 | 1.98E-06 |
| Sleep Duration | Muscle_Skeletal | GLTP | -4.73 | 2.27E-06 |
| Sleep Duration | Nerve_Tibial | GLTP | -4.76 | 1.89E-06 |
| Sleep Duration | Ovary | GLTP | -4.75 | 2.02E-06 |
| Sleep Duration | Pancreas | GLTP | -4.76 | 1.98E-06 |
| Sleep Duration | Prostate | GLTP | 4.73 | 2.26E-06 |
| Sleep Duration | Skin_Not_Sun_Exposed_Suprapubic | GLTP | -4.75 | 2.03E-06 |
| Sleep Duration | Small_Intestine_Terminal_Ileum | GLTP | 4.77 | 1.88E-06 |
| Sleep Duration | Spleen | GLTP | 4.76 | 1.94E-06 |
| Sleep Duration | Stomach | GLTP | -4.77 | 1.89E-06 |
| Sleep Duration | Testis | GLTP | 4.75 | 2.06E-06 |
| Sleep Duration | Uterus | GLTP | 4.73 | 2.28E-06 |
| Sleep Duration | Whole_Blood | GLTP | 4.67 | 2.94E-06 |
| Sleep Duration | Lung | KIFC3 | 4.65 | 3.28E-06 |
| Sleep Start | Artery_Coronary | ABCD2 | 4.71 | 2.53E-06 |
| Sleep Start | Brain_Anterior_cingulate_cortex_BA24 | ABCD2 | -4.61 | 3.94E-06 |
| Sleep Start | Adipose_Visceral_Omentum | ABCD2 | 4.76 | 1.94E-06 |
| Sleep Start | Brain_Cortex | ABCD2 | 4.62 | 3.90E-06 |
| Sleep Start | Brain_Cerebellum | ABCD2 | -4.78 | 1.79E-06 |
| Sleep Start | Colon_Sigmoid | ABCD2 | 4.61 | 4.11E-06 |
| Sleep Start | Adipose_Subcutaneous | ABCD2 | 4.75 | 2.03E-06 |
| Sleep Start | Colon_Transverse | ABCD2 | 4.67 | 3.02E-06 |
| Sleep Start | Whole_Blood | ABCD2 | -4.58 | 4.55E-06 |
| Sleep Start | Small_Intestine_Terminal_Ileum | ALG10B | 4.94 | 7.99E-07 |
| Sleep Start | Whole_Blood | ALG10B | -4.75 | 1.99E-06 |
| Sleep Start | Adipose_Subcutaneous | CCDC181 | -4.81 | 1.49E-06 |
| Sleep Start | Lung | CCDC87 | -4.63 | 3.66E-06 |
| Sleep Start | Brain_Anterior_cingulate_cortex_BA24 | CEP128 | -4.62 | 3.85E-06 |
| Sleep Start | Prostate | FSIP2 | -4.65 | 3.28E-06 |
| Sleep Start | Nerve_Tibial | KIAA0391 | -4.65 | 3.31E-06 |
| Sleep Start | Brain_Cerebellum | LIMS1 | 4.90 | 9.76E-07 |
| Sleep Start | Brain_Nucleus_accumbens_basal_ganglia | LIMS1 | 5.02 | 5.18E-07 |
| Sleep Start | Testis | LIMS1 | -4.81 | 1.50E-06 |
| Sleep Start | Brain_Frontal_Cortex_BA9 | LIMS1 | 4.95 | 7.61E-07 |
| Sleep Start | Esophagus_Mucosa | LRFN4 | 4.80 | 1.62E-06 |
| Sleep Start | Brain_Anterior_cingulate_cortex_BA24 | LRFN4 | 4.63 | 3.59E-06 |
| Sleep Start | Adrenal_Gland | LRFN4 | 4.65 | 3.26E-06 |
| Sleep Start | Artery_Coronary | LRFN4 | 4.62 | 3.76E-06 |
| Sleep Start | Colon_Sigmoid | LRFN4 | 4.64 | 3.55E-06 |
| Sleep Start | Ovary | LRFN4 | 4.79 | 1.65E-06 |
| Sleep Start | Uterus | MYO18A | -5.05 | 4.39E-07 |
| Sleep Start | Brain_Cortex | STX17 | -4.65 | 3.37E-06 |
| Sleep Start | Colon_Transverse | TREH | 7.23 | 4.68E-13 |
| Sleep Start | Stomach | TREH | 5.76 | 8.54E-09 |
| Sleep Start | Esophagus_Mucosa | TREH | 5.53 | 3.15E-08 |
| Sleep End | Whole_Blood | HILPDA | -4.66 | 3.19E-06 |
| Sleep End | Brain_Cerebellum | JARID2 | -5.13 | 2.95E-07 |
| Sleep End | Artery_Coronary | SEPN1 | -4.69 | 2.73E-06 |
| Sleep End | Brain_Cortex | SEPN1 | -4.61 | 3.94E-06 |
| 1-day Periodicity | Brain_Cerebellar_Hemisphere | C1orf162 | 4.91 | 9.06E-07 |
| 1-day Periodicity | Liver | C1orf162 | -4.94 | 7.97E-07 |
| 1-day Periodicity | Adipose_Subcutaneous | L3MBTL2 | -5.03 | 4.81E-07 |
| 1-day Periodicity | Breast_Mammary_Tissue | L3MBTL2 | -4.77 | 1.82E-06 |
| 1-day Periodicity | Muscle_Skeletal | L3MBTL2 | -4.83 | 1.38E-06 |
| 1-day Periodicity | Muscle_Skeletal | SPARC | -5.28 | 1.30E-07 |
| 1-day Periodicity | Artery_Tibial | ZFAND2B | 4.73 | 2.26E-06 |
| 1-day Periodicity | Cells_EBV-transformed_lymphocytes | ZNF79 | -4.63 | 3.72E-06 |
| 1/2-day Periodicity | Esophagus_Mucosa | CPZ | -4.78 | 1.79E-06 |
| 1/2-day Periodicity | Pancreas | DCTPP1 | 4.72 | 2.38E-06 |
| 1/2-day Periodicity | Esophagus_Mucosa | PCDH9 | 4.61 | 3.98E-06 |
| 1/2-day Periodicity | Muscle_Skeletal | PCDH9 | 4.61 | 3.98E-06 |
| 1/2-day Periodicity | Prostate | PCDH9 | -4.61 | 3.98E-06 |
| 1/2-day Periodicity | Vagina | PCDH9 | 4.61 | 3.98E-06 |
| 1/2-day Periodicity | Brain_Hypothalamus | PRSS36 | 4.65 | 3.39E-06 |
| 1/2-day Periodicity | Brain_Putamen_basal_ganglia | STARD3 | -4.89 | 1.02E-06 |
| 1/3-day Periodicity | Brain_Anterior_cingulate_cortex_BA24 | PDE7B | 4.63 | 3.59E-06 |
| 1/3-day Periodicity | Muscle_Skeletal | STATH | 4.97 | 6.74E-07 |

**Table S5.** Cross-tissue gene-trait association test results from tissue enrichment analysis using UTMOST.

| Phenotype | gene | test_score | p_value |
| --- | --- | --- | --- |
| Mean Activity Levels During Sleep | CA7 | 13.14 | 2.53E-06 |
| Mean Activity Levels During Sleep | LYVE1 | 13.80 | 2.48E-06 |
| Mean Activity Levels During Sleep | PIAS2 | 15.82 | 6.13E-08 |
| Activity Variability During Sleep | FTL | 21.53 | 1.15E-09 |
| Mean Activity Levels During Wake | CA7 | 23.03 | 1.28E-10 |
| Mean Activity Levels During Wake | L3MBTL2 | 12.32 | 2.16E-06 |
| Mean Activity Levels During Wake | TRAF3 | 14.66 | 3.35E-07 |
| Activity Variability During Wake | CA7 | 27.10 | 5.19E-12 |
| Activity Variability During Wake | DUSP15 | 12.25 | 1.64E-06 |
| Activity Variability During Wake | ELMOD2 | 43.91 | 2.91E-11 |
| Activity Variability During Wake | POM121L7 | 13.40 | 1.96E-06 |
| Activity Variability During Wake | SLC7A4 | 12.85 | 2.20E-06 |
| Sleep Duration | DYNC1LI1 | 17.77 | 5.76E-08 |
| Sleep Duration | ELMOD2 | 59.88 | 2.91E-11 |
| Sleep Duration | GDPD5 | 15.22 | 1.56E-07 |
| Sleep Duration | GLTP | 14.21 | 1.11E-06 |
| Sleep Start | ABCD2 | 150.41 | 8.38E-07 |
| Sleep Start | DCLK2 | 13.67 | 7.72E-07 |
| Sleep Start | DYNC1LI1 | 49.66 | 2.91E-11 |
| Sleep Start | KIDINS220 | 17.58 | 1.40E-08 |
| Sleep Start | PPP2R2C | 13.14 | 1.16E-06 |
| Sleep Start | TREH | 23.76 | 2.01E-11 |
| Sleep End | ELMOD2 | 54.35 | 2.91E-11 |
| Sleep End | GRIA1 | 13.33 | 1.44E-06 |
| Sleep End | VCAM1 | 35.73 | 2.91E-11 |
| 1-day Periodicity | C1orf162 | 12.84 | 1.90E-06 |
| 1/2-day Periodicity | ALS2 | 11.95 | 2.28E-06 |
| 1/2-day Periodicity | ALS2CR11 | 19.56 | 1.64E-09 |
| 1/2-day Periodicity | CPZ | 14.81 | 2.12E-07 |
| 1/2-day Periodicity | DPP9 | 16.21 | 2.84E-08 |
| 1/2-day Periodicity | DYNC1LI1 | 21.36 | 1.63E-09 |
| 1/2-day Periodicity | ELMOD2 | 20.29 | 4.83E-09 |
| 1/3-day Periodicity | DYNC1LI1 | 27.16 | 3.36E-11 |
| 1/3-day Periodicity | ELMOD2 | 23.54 | 1.98E-10 |

**Table S6.** Two-sample Mendelian Randomization analysis for the strength of circadian rhythm using GWAS summary statistics from the GIANT study.

| Exposure | Outcome | Outcome units | Method | N SNPs | beta | SE | P Value |
| --- | --- | --- | --- | --- | --- | --- | --- |
| Strength of Circadian Rhythm | Body mass index (BMI) | SD (Kg/m<sup>2</sup>) | Inverse variance weighted | 11 | -0.00050 | 0.00014 | 3.42E-04 |
| Strength of Circadian Rhythm | Body mass index (BMI) | SD (Kg/m<sup>2</sup>) | Weighted median | 11 | -0.00067 | 0.00010 | 1.81E-11 |
| Strength of Circadian Rhythm | Body mass index (BMI) | SD (Kg/m<sup>2</sup>) | Weighted mode | 11 | -0.00071 | 0.00011 | 7.93E-05 |
| Strength of Circadian Rhythm | Body mass index (BMI) | SD (Kg/m<sup>2</sup>) | MR Egger | 11 | -0.00152 | 0.00103 | 0.174 |

